# A yeast-based reverse genetics system to generate HCoV-OC43 reporter viruses encoding an eighth sgRNA

**DOI:** 10.1101/2024.09.23.614401

**Authors:** Brett A. Duguay, Trinity H. Tooley, Eric S. Pringle, John R. Rohde, Craig McCormick

## Abstract

Coronaviruses have large, positive-sense single-stranded RNA genomes that challenge conventional strategies for mutagenesis. Here, we report the development of a new reverse genetics system for the endemic human coronavirus (HCoV) OC43 that utilizes transformation-associated recombination (TAR) to assemble complete viral genomes from dsDNA genome fragments via homologous recombination in *Saccharomyces cerevisiae*. Following cDNA synthesis from HCoV-OC43 viral RNA, we used TAR to capture fragments of the HCoV-OC43 genome to store as sequence-validated dsDNA parts. We performed combinatorial assembly in yeast to obtain an intact dsDNA copy of the HCoV-OC43 genome sufficient to launch viral replication upon introduction into human cells, yielding the yeast assembled OC43^YA^ virus. We also expanded the OC43^YA^ genome by inserting an eighth body transcription regulatory sequence (B-TRS) and an mClover3-H2B reporter gene between the *M* and *N* genes, designed to allow the reporter protein to be translated from its own subgenomic mRNA. We thoroughly evaluated OC43^YA^ and the OC43-mClo^YA^ reporter virus, and demonstrated comparable viral gene expression, fitness in cell culture, and susceptibility to antivirals, compared to their natural progenitor. In summary, this new HCoV-OC43 reverse genetics system provides a modular platform for mutagenesis and combinatorial assembly of HCoV-OC43 genomes, and demonstrates the feasibility of expanding the genome while avoiding disruption of native coding sequences.

**IMPORTANCE:** Human coronavirus OC43 (HCoV-OC43) is an endemic human coronavirus that typically causes relatively mild respiratory illnesses and displays seasonal patterns of infection. We developed a new system to assemble DNA copies of HCoV-OC43 genomes and generate recombinant viruses for research purposes. This system uses yeast, first to capture segments of DNA encompassing the entire RNA-based viral genome, and then to stitch them together into complete DNA genome copies that can be amplified in bacteria and introduced into human cells to initiate an infectious cycle, ultimately yielding recombinant viruses with comparable properties to their natural progenitors. We also devised a strategy to expand the viral genome, adding a gene for a reporter protein encoded by an additional eighth subgenomic mRNA. This yeast-based genome assembly system provides a modular platform for rapid mutagenesis and combinatorial assembly of HCoV-OC43 genomes and demonstrates the feasibility of expanding the genome.

## INTRODUCTION

Coronaviruses, from the order Nidovirales, are positive-sense, single-stranded RNA ((+)ssRNA) viruses that possess some of the largest (12-41 kb) monopartite RNA genomes (1–3). These viruses encode a large polyprotein from *ORF1ab* encompassing the first two-thirds of the genome, which is translated using ribosomal frameshifting RNA elements (FSEs) into polyproteins pp1a and pp1ab (Fig. 1A and (4)). These polyproteins are proteolytically processed by the viral papain-like protease (PLpro) and the main protease (Mpro) into 16 non-structural proteins (Nsps) with roles in host subversion, replication compartment formation, and viral RNA production and modification (5). The remaining 3’ third of coronavirus genomes encode accessory and structural proteins. These proteins are translated from a nested set of subgenomic mRNAs (sg-mRNAs) produced through discontinuous transcription facilitated by transcription regulatory sequences (TRSs), with TRS-L positioned at the 5’ end of the genome and numerous TRS-B elements dispersed near the 3’ end of the genome, in regions that aid production of diverse sg-mRNAs (Fig. 1A). These regulatory sequences direct template switching by the viral RNA-dependent RNA polymerase during negative-sense subgenomic RNA ((-)sgRNA) transcription to produce templates for subsequent viral sg-mRNA synthesis (6–8).

**Figure 1.**
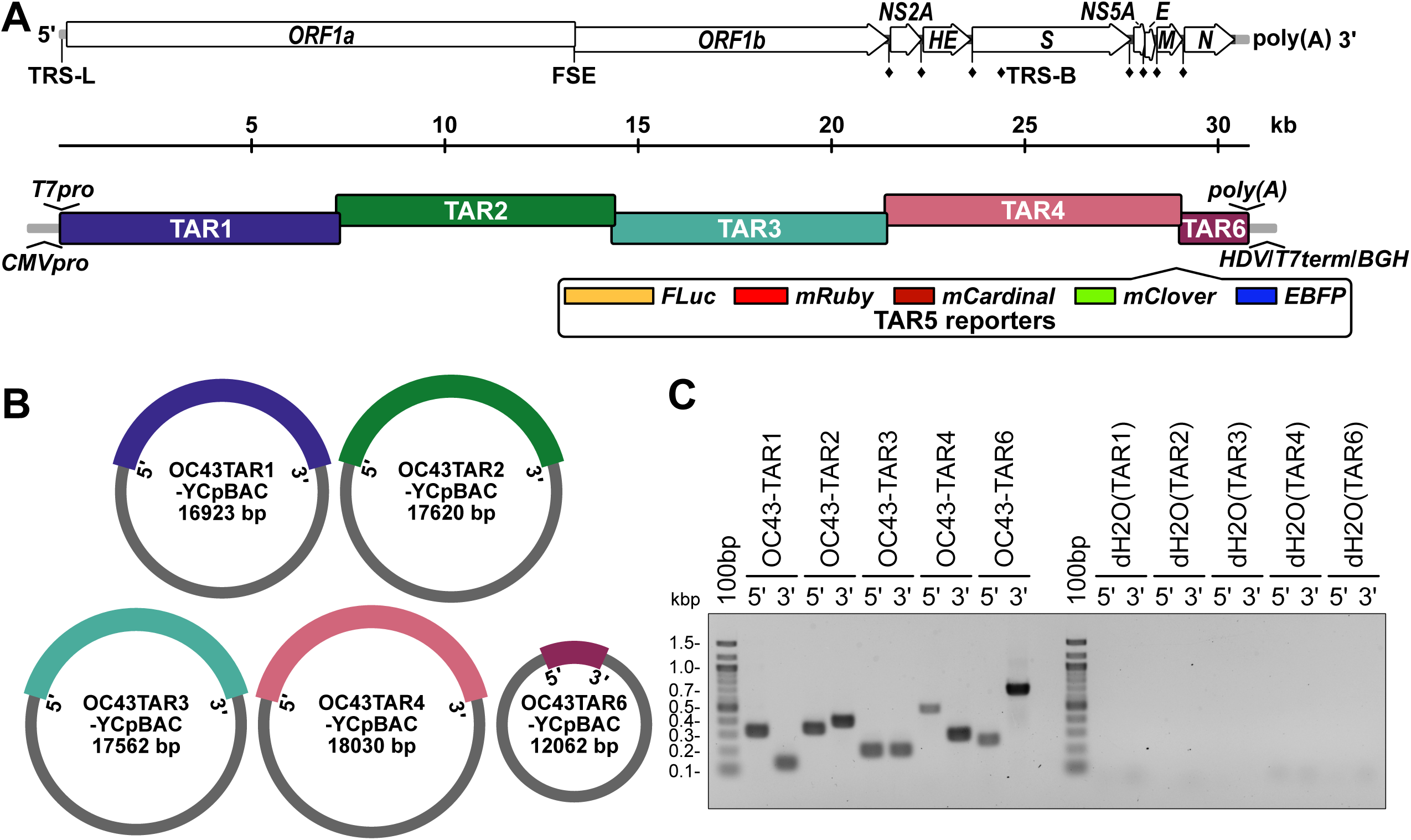
Design of HCoV-OC43 TAR plasmids for yeast assembly of viral sequences. **(A)** A schematic of the HCoV-OC43 genome including annotation of the leader transcription regulatory sequence (TRS-L), ribosomal frameshifting RNA element (FSE), and body TRSs (TRS-B, **♦**) (top) and the orientations of the six, overlapping coronavirus TAR fragments (bottom). Inserted regulatory sequences are indicated at the 5’ (*T7pro*, T7 polymerase promoter; *CMVpro*, cytomegalovirus promoter) and 3’ (*poly(A)* tail; *HDV*, hepatitis delta virus ribozyme; *T7term*, T7 terminator; *BGH*, bovine growth hormone polyadenylation signal) ends of the genome. Five different optional reporter genes (TAR5 reporters) are designed to be inserted between the *M* and *N* genes. **(B)** Illustration of the circular yeast centromeric plasmid/bacterial artificial chromosome (YCpBAC) vectors generated through homologous recombination in yeast with the corresponding HCoV-OC43 TAR sequences. HCoV-OC43 sequences are colour-coded to match the diagram in panel A. YCpBAC sequences are shown in grey. **(C)** Plasmids containing the indicated HCoV-OC43 TAR fragments were subjected to PCR using primer sets designed to amplify the 5’ or 3’ junctions between the HCoV-OC43 TAR fragments and the YCpBAC sequences. PCR water controls (dH2O) are shown to the right of the panel. Primer pairs used for screening are listed in Table 2.

The development of reverse genetics systems accelerated study of coronaviruses. These systems work on the premise of regenerating an infectious genomic RNA (gRNA) from a plasmid assembled using complementary DNA (cDNA) fragments. Due to the challenge of working with such large genomes, multiple strategies have been adopted to reassemble full-length coronavirus genomic sequences including: RNA recombination (9–11), restriction digestion and *in vitro* ligation (12–32), lambda red-mediated recombination in bacteria (33, 34), Gibson assembly (35), circular polymerase extension reaction (CPER) (36–40), Golden Gate assembly (41), infectious subgenomic amplicons (42, 43), and transformation-associated recombination (TAR) in yeast (44–46).

Based on our experience using the human coronavirus OC43 (HCoV-OC43) as a biosafety level 2 model of betacoronavirus infection, we set out to develop a yeast-based genome assembly and mutagenesis platform of HCoV-OC43 to support antiviral drug discovery and host-pathogen interaction research. The TAR method uses the highly efficient *Saccharomyces cerevisiae* homologous recombination system to join numerous transformed DNA fragments with ≥40 bp of DNA homology into yeast centromeric plasmids (YCp) (47). It has been used to assemble genomes of large DNA viruses including herpes simplex virus type 1, human cytomegalovirus (HCMV), Kaposi’s sarcoma-associated herpesvirus (48–50) and to generate cDNA copies of RNA viruses such as: swine enteric alphacoronavirus, mouse hepatitis virus (MHV), Middle East respiratory syndrome coronavirus, severe acute respiratory syndrome coronavirus 2 (SARS-CoV-2), Japanese encephalitis virus, porcine epidemic diarrhea virus, and feline coronavirus (44–46, 51, 52).

Prior to beginning our work on HCoV-OC43, only one bacterial artificial chromosome (BAC)-based reverse genetic platform for this virus had been developed (18). Earlier this year, two additional HCoV-OC43 mutagenesis systems were published, both of which generated *NS2A* substitution mutants (32, 40). Here we describe our work applying synthetic biology techniques to develop an HCoV-OC43 reverse genetics system culminating in the generation of <u>y</u>east <u>a</u>ssembled wild-type (OC43^YA^) and mClover3-H2B reporter (OC43-mClo^YA^) viruses. The viruses we assembled and rescued replicated comparably to a wild-type strain of HCoV-OC43. Importantly, our updated reporter virus platform did not require the mutation of any viral protein. By expressing the mClover3 variant of GFP (53) fused to histone H2B, this reporter provides improved brightness and exclusively nuclear localization compared to EGFP, making it well suited for live cell imaging, immunofluorescence, automated image analysis, and flow cytometric analyses of viral replication. We have made plasmids containing individual cDNA fragments or partially assembled cDNAs of the viral genome to provide some modularity to the assembly process and to maximize flexibility for performing mutagenesis. Generation of the OC43-mClo^YA^ reporter virus demonstrates that HCoV-OC43 tolerates both a 1223 base insertion in the intergenic region between the *M* and *N* genes and the addition of an eighth TRS-B. The OC43-mClo^YA^ virus is the first example of an HCoV-OC43 reporter virus built without the deletion or mutation of viral coding sequences.

## RESULTS

Viral reverse genetics systems have greatly impacted the understanding of viral biology and can accelerate the study of antiviral interventions by producing reporter viruses to rapidly track infectivity. The first BAC-based, HCoV-OC43 reverse genetics system was developed using overlap PCR products and restriction cloning (18), which led to the creation of luciferase reporter viruses deficient in *NS2A* or *NS5A* (54). More recently, HCoV-OC43 reverse genetics systems using *in vitro* ligation or CPER were used to generate Δ*NS2A*::*nanoluciferase* or ORF1b-EGFP fusion reporter viruses, respectively (32, 40). Previously, work with another betacoronavirus, MHV, demonstrated the ability to insert luciferase genes preceded by independent TRSs to drive reporter gene expression; where reporter gene insertions proximal to the 3’ end of the viral genome resulted in the most robust expression (10). We reasoned that HCoV-OC43 viruses encoding fluorescent protein reporter genes could be generated using a similar strategy. By introducing an additional TRS into the viral genome, this strategy eliminates the need for fusion proteins or the deletion of viral accessory genes in recombinant HCoV-OC43 strains.

### Design, capture, and assembly of HCoV-OC43 cDNA plasmids

We developed an updated reverse genetics system for HCoV-OC43 that employs efficient homologous recombination in *S. cerevisiae* to assemble viral cDNA fragments. This method of TAR cloning has been applied to cloning complete human genes, yeast chromosomes, and to the assembly of entire bacterial genomes through accurate, stable, recombination-mediated DNA assembly (55–58). Our strategy divided the viral genome into five fragments: Three ∼7.2 kb portions of the *ORF1ab* genes (TAR1, 2, and 3), a fourth 7.4 kb portion encoding the *NS2* to *M* genes (TAR4), and a short fifth fragment (1.6 kb) containing the *N* gene (TAR6) (Fig. 1A).

We employed two strategies for the rescue of infectious HCoV-OC43, T7 RNA polymerase-directed gRNA production or CMV promoter-driven *in vivo* rescue, which required the insertion of T7 RNA polymerase promoter (*T7pro*) or CMV/RNA polymerase II promoter (*CMVpro*) sequences directly 5’ to the HCoV-OC43 5’ untranslated region (5’UTR) in TAR1 (Fig. 1A). Likewise, essential regulatory sequences were inserted downstream of *N*, including a synthetic stretch of 34 adenosine monophosphates (*poly(A)*) for RNA stability and a hepatitis delta ribozyme self-cleaving RNA sequence (*HDV*; (59, 60)) to generate an authentic gRNA 3’ end (Fig. 1A). Downstream of the ribozyme sequence were a T7 RNA polymerase terminator (*T7term*), to use with the *T7pro* system, and a bovine growth hormone polyadenylation signal (*BGH*), for use with the *CMVpro* system (Fig. 1A). First, HCoV-OC43 gRNA was converted to cDNA and PCR amplified. These PCR amplicons where then co-transformed into yeast along with a linearized yeast centromeric plasmid/bacterial artificial chromosome (YCpBAC) plasmid containing unique 5’ and 3’ ends homologous to the various HCoV-OC43 TAR fragments (Fig. 1B, Table 1). The resulting yeast clones were screened for the 5’ and 3’ DNA junctions between the HCoV-OC43 and YCpBAC sequences and yielded banding patterns indicative of correct fragment capture into the YCpBACs (Fig. 1C, Table 2).

**Table 1:**
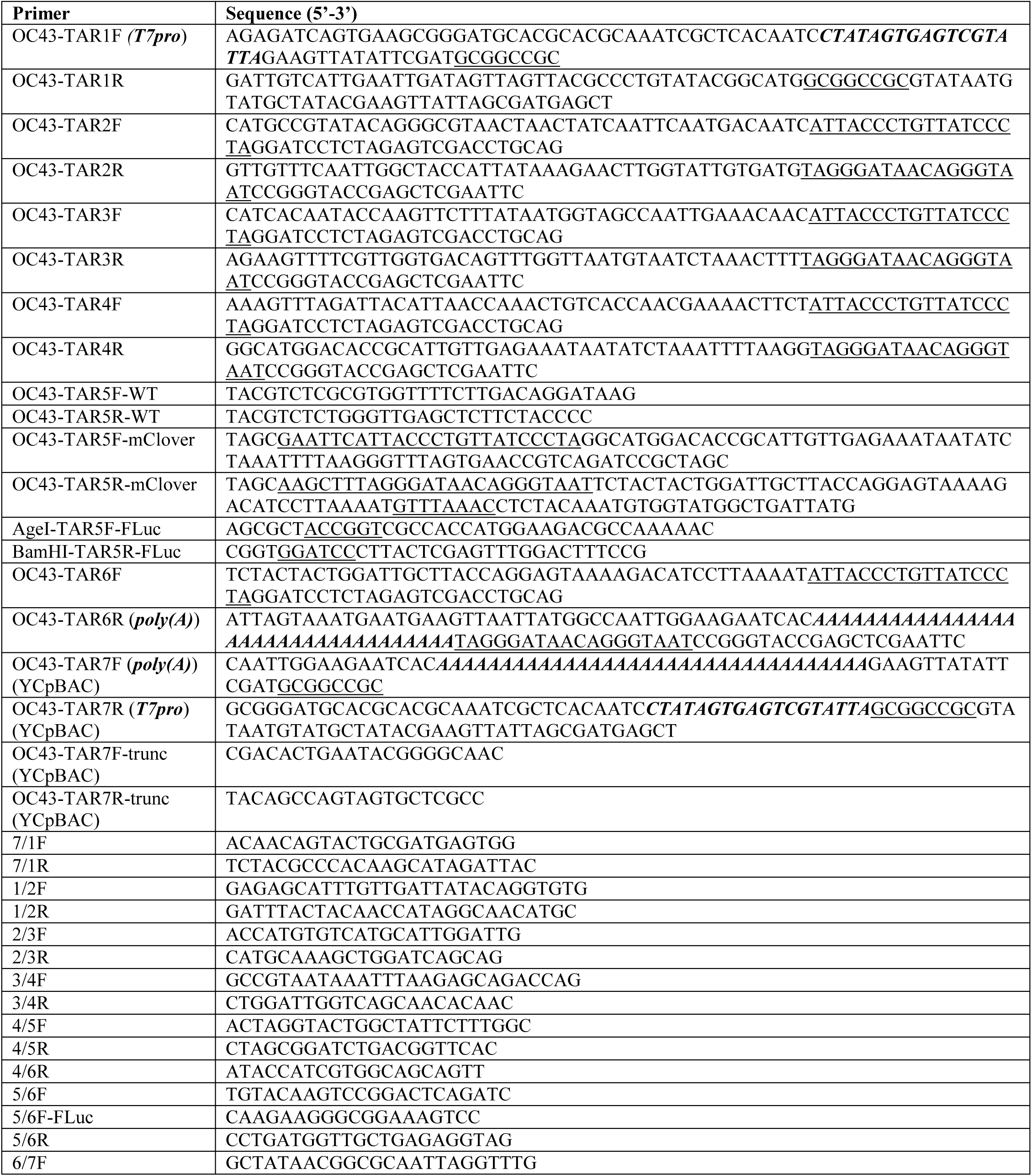
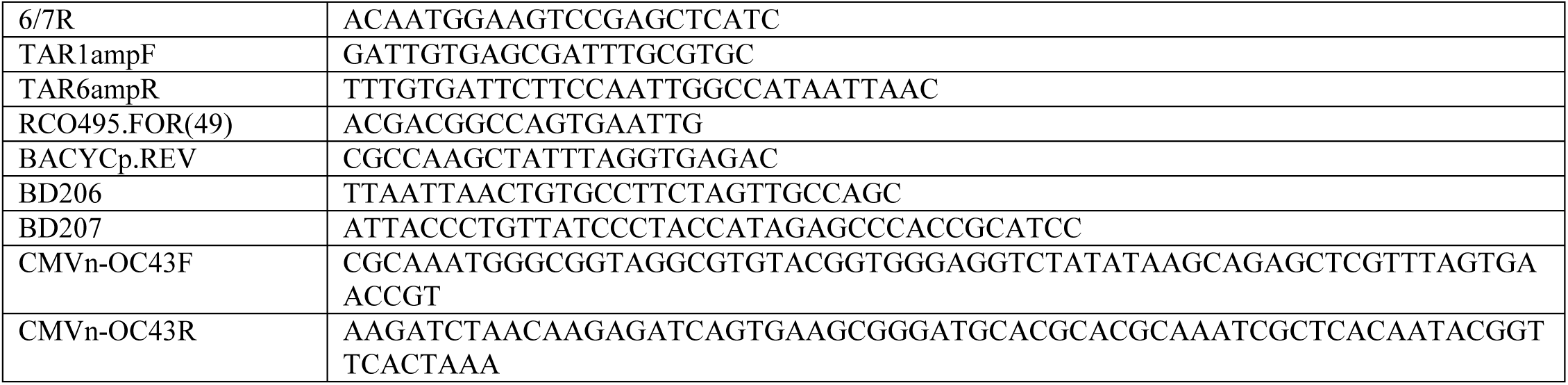
Oligonucleotides for TAR cloning and screening.

HCoV-OC43 sequences were reassembled into *Orf1ab* (TAR123) or *NS2A*-to-*N* (TAR456) containing YCpBACs to facilitate subsequent full-length HCoV-OC43 sequence assemblies (Figs. 2A and 2B). The three TAR plasmids encoding fragments of *Orf1ab* were linearized and co-transformed with a YCpBAC (TAR7, in grey) sequence to facilitate assembly of OC43TAR123-YCpBAC in yeast (Fig. 2A). The correct junction amplicons were obtained during screening for the junctions 7/1, 1/2, 2/3, and 3/7 (Fig. 2C, Table 2), indicating the correct assembly of these sequences. Likewise, the three TAR plasmids encoding *NS2A*, *HE*, *S*, *NS5A*, *E*, and *M* (TAR4), the reporter genes *mClover3-H2B* (mClover3 fused to histone H2B), *mRuby3-H2B* (mRuby3 fused to histone H2B), *mCardinal*, *EBFP2*, or firefly *luciferase* (*FLuc*) (TAR5), and *N* (TAR6) were linearized and co-transformed into yeast with a YCpBAC (TAR7, in grey) to assemble the OC43TAR456-YCpBAC plasmids (Fig. 2B). This design provided the flexibility to insert various reporter gene sequences, driven from the authentic *N* TRS, into the region between *M* and *N*. Since the authentic TRS for *N* now drives reporter gene sgRNA synthesis, the expression of *N* was maintained through the insertion of a duplicated 23 base TRS-containing element upstream of the *N* start codon (TRS*, Fig. 2D). The correct junction amplicons were obtained during screening for the junctions 7/4, 4/5, 5/6, 5F/6 (FLuc only), and 6/7 (Fig. 2E and 2F, Table 2), indicating the correct assembly of the reporter plasmids. Assembly of a wild-type *NS2A*-*N* sequence lacking a reporter gene was completed using a 297 bp TAR5 PCR amplicon. To provide the flexibility to insert sequences between *M* (TAR4) and *N* (TAR6), these TAR fragments have minimal overlap, thus requiring the short PCR amplicon for assembly. The anticipated DNA junction amplicons (7/4, 4/6, and 6/7) were obtained for this wild-type plasmid (Fig. 2G, Table 2).

**Figure 2.**
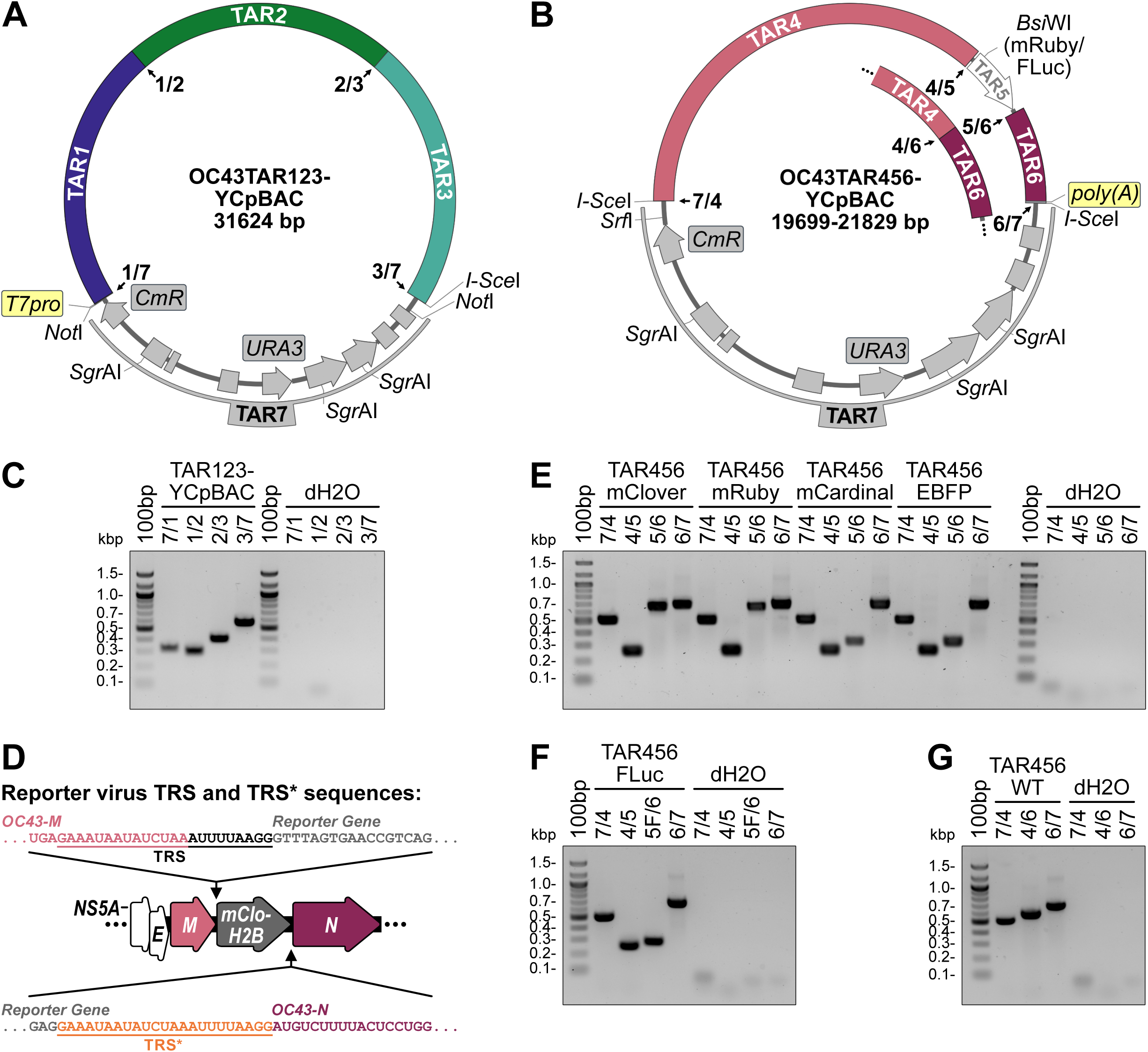
Yeast assembly of plasmids encoding HCoV-OC43 *Orf1ab* and *NS2A-to-N*. Diagrams of the OC43TAR123-YCpBAC plasmid containing HCoV-OC43 TAR fragments 1, 2, and 3 encoding *Orf1ab* **(A)** and OC43TAR456-YCpBACs containing HCoV-OC43 TAR fragments 4, 5 (indicated in white in the plasmid map), and 6 or fragments (for reporter virus assembly) or 4 and 6 (for wild-type (WT) virus assembly; inset) encoding *NS2A* to *N* +/− reporter genes **(B)**. HCoV-OC43 sequences are colour-coded to match the diagram in Fig. 1. YCpBAC sequences (TAR7) are shown in grey. Numbered junctions detected by PCR (black arrows) and restriction sites used for TAR cloning are indicated. **(C)** Following assembly in yeast, the TAR123-YCpBAC was subjected to PCR using primer sets designed to amplify junctions 7/1, 1/2, 2/3, and 3/7 with water controls (dH2O) to the right of the panel. **(D)** Arrangement of the *M*-*N* locus in reporter virus genomes with the authentic *N* TRS used to drive reporter gene sgRNA synthesis and an inserted copy of the *N* TRS (TRS*, sequence in orange) for *N* sgRNA synthesis indicated. **(E)** Following assembly in yeast, TAR456-YCpBACs were subjected to PCR using primer sets designed to amplify junctions 7/4, 4/5 (present when flanking a reporter gene), 5/6 (for fluorescence reporter genes) and 6/7 with dH2O controls to the right of the panel. **(F)** PCR screening of assembled TAR456-FLuc-YCpBAC with screening for junctions: 7/4, 4/5, 5F/6 (for the FLuc reporter gene), and 6/7 with dH2O controls to the right of the panel. **(G)** PCR screening of assembled TAR456-WT-YCpBAC with screening for junctions: 7/4, 4/6, and 6/7 with dH2O controls to the right of the panel. Primer pairs used for screening are listed in Table 2. Abbreviations: 100bp, 100 bp ladder; bp/kbp, base/kilobase pairs; *CmR*, chloramphenicol resistance gene; EBFP, enhanced blue fluorescent protein; FLuc, firefly luciferase; *poly(A)*, encoded A_34_ sequence; *T7pro*, T7 polymerase promoter; TRS, transcription regulatory sequence; *URA3*, orotidine 5’-phosphate decarboxylase gene; WT, wild-type.

**Table 2:** Screening amplicon sizes following YCpBAC assembly.

| **Amplicon (primer pair)** | **Size (bp)** | **Amplicon (primer pair)** | **Size (bp)** |
| --- | --- | --- | --- |
| OC43-TAR1 5' (7/1F+7/1R) | 321 | OC43-TAR1 3' (1/2F+6/7R) | 138 |
| OC43-TAR2 5' (BACYcp.REV+1/2R) | 349 | OC43-TAR2 3' (2/3F+ RCO495.FOR) | 396 |
| OC43-TAR3 5' (BACYcp.REV+2/3R) | 212 | OC43-TAR3 3' (3/4F+ RCO495.FOR) | 212 |
| OC43-TAR4 5' (BACYcp.REV+3/4R) | 508 | OC43-TAR4 3' (4/5F+ RCO495.FOR) | 301 |
| OC43-TAR6 5' (BACYcp.REV+5/6R) | 264 | OC43-TAR6 3' (6/7F+ RCO495.FOR) | 693 |
| 7/1 (*T7pro*) (7/1F+7/1R) | 321 | 7/1 (*CMVpro*) (7/1F+7/1R) | 879 |
| 1/2 (1/2F+1/2R) | 294 | 2/3 (2/3F+2/3R) | 397 |
| 3/4 (3/4F+3/4R) | 509 | 3/7 (3/4F+6/7R) | 571 |
| 4/5 (4/5F+4/5R) | 249 | 4/6 (4/5F+4/6R) | 595 |
| 5/6 (mClover) (5/6F+5/6R) | 660 | 5/6 (mRuby) (5/6F+5/6R) | 660 |
| 5/6 (mCardinal) (5/6F+5/6R) | 312 | 5/6 (EBFP) (5/6F+5/6R) | 312 |
| 5F/6 (FLuc) (5/6F-FLuc+5/6R) | 276 | 6/7 (*-Ribo*) (6/7F+ RCO495.FOR) | 693 |
| 6/7* (*+Ribo*) (6/7F+6/7R) | 1207 | 7/4 (BACYcp.REV+3/4R) | 508 |

To assemble the full-length HCoV-OC43 viral sequences, we co-transformed linearized OC43TAR123-YCpBAC, OC43TAR456-YCpBAC, and amplified YCpBAC sequences into yeast (Fig. 3A). Performing full-length assemblies with three fragments consistently yielded more correct transformants than seven-part assemblies (data not shown). During the assembly of HCMV genomes via TAR, similar three-part, “half-genome” assemblies were also more successful than 17 fragment assemblies (48). Both RNA-based and DNA-based approaches have been used to successfully rescue coronaviruses from plasmids. This includes *in vitro* transcription and capping of a genome length transcript for delivery into cells, *T7pro* driven transcription with ectopically expressed T7 RNA polymerase in plasmid transfected cells, as well as *CMVpro* driven transcription in plasmid transfected cells. We initially designed our system to generate full-length mRNA from a *T7pro* with a hepatitis delta virus ribozyme (*Ribo*) inserted following the encoded poly(A) sequence (Fig. 3A, top line). However, these constructs failed to yield infectious virus from transfected *in vitro* synthesized and capped mRNA or from a plasmid-based, co-transfection with T7 RNA polymerase and vaccina virus D1R capping enzyme plasmids (data not shown). Therefore, we transitioned to a DNA-based rescue system using a CMV promoter (*CMVpro*) upstream of the HCoV-OC43 5’UTR and a bovine growth hormone polyadenylation signal (*BGH*) downstream of the *Ribo* sequence which were inserted into the OC43-mClover-Ribo-YCpBAC using homologous recombination in yeast (Fig. 3A, bottom line). Our first *CMVpro*-driven constructs containing *CMVpro* and *T7pro* sequences upstream of the 5’UTR similarly did not yield infectious virus (data not shown). Ultimately, the constructs that yielded infectious HCoV-OC43 utilized a CMV enhancer/promoter sequence positioned 15 bp upstream of the 5’UTR (*CMVn*) as designed by St-Jean *et al*. during the generation of an HCoV-OC43 BAC (18). The final full-length assemblies of CMVn-OC43-mClover-Ribo-BGH-YCpBAC (Fig. 3B-C) and CMVn-OC43-WT-Ribo-BGH-YCpBAC (wild-type sequence; Fig. 3D-E) yielded the correct DNA junction amplicons (Table 2) as shown in Fig. 3C and Fig. 3E.

**Figure 3:**
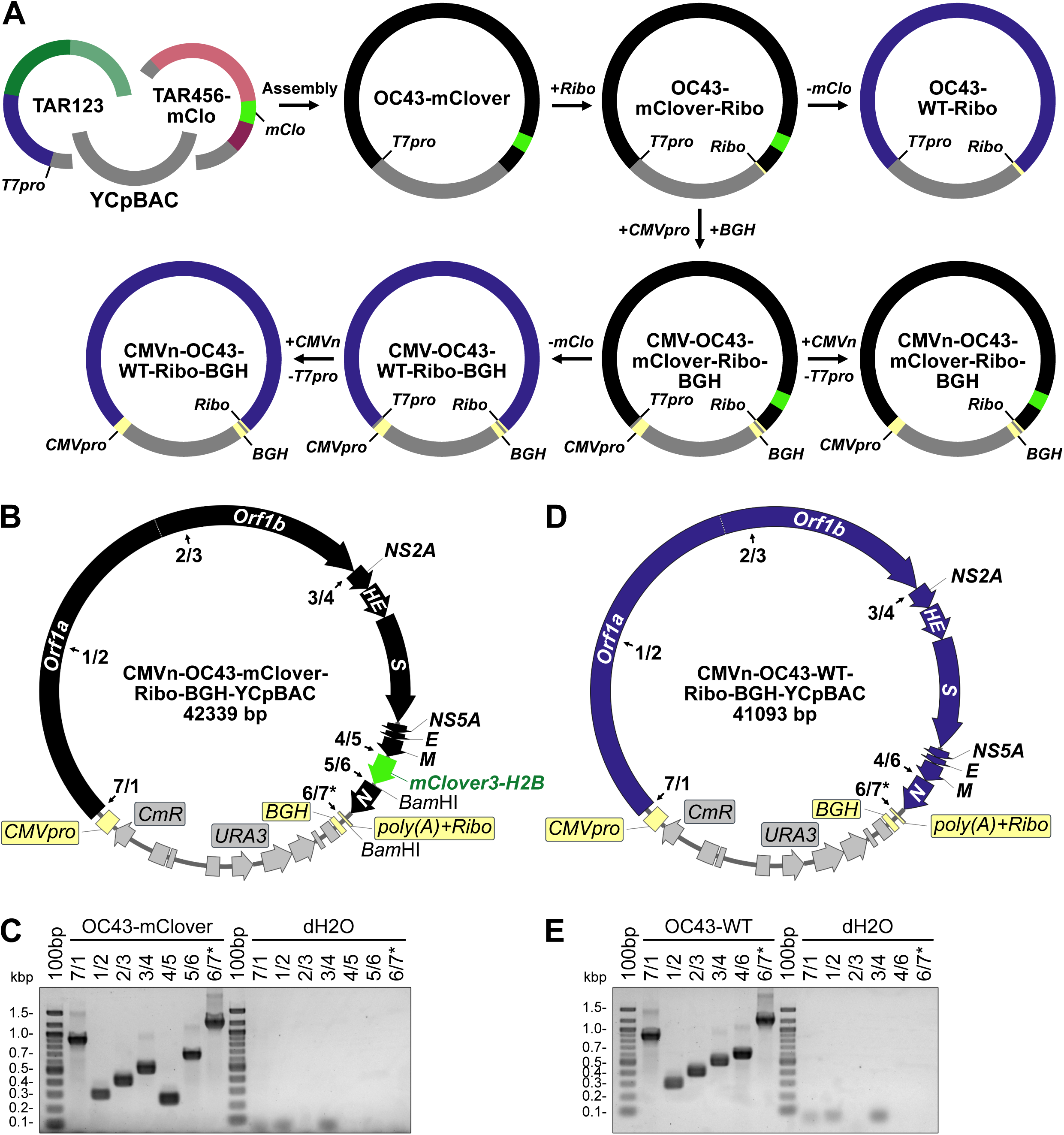
Yeast assembly of full-length HCoV-OC43 reporter and wild-type virus plasmids. **(A)** Assembly of a full-length HCoV-OC43-mClover viral sequences (black) was performed by sequential assembly steps in yeast. First, restriction digested OC43TAR123-YCpBAC and OC43TAR456-mClover plasmids were assembled, followed by the sequential insertion of hepatitis delta virus ribozyme (*Ribo*), bovine growth hormone poly(A) signal (*BGH*), and CMV promoter (*CMVpro*) sequences (light yellow). Assembly of full-length HCoV-OC43-WT viral sequences (blue) was performed by removing the mClover3-H2B (*mClo*, light green) sequence from previously assembled plasmids. **(B)** Plasmid map for the CMVn-OC43-mClover-Ribo-BGH-YCpBAC used for virus rescue. Numbered junctions detected by PCR (black arrows) and restriction sites used for TAR cloning are indicated. **(C)** PCR screening of assembled junctions 7/1, 1/2, 2/3, 3/4, 4/5, 5/6, and 6/7* with water controls (dH2O) to the right of the panel. **(D)** Plasmid map for the CMVn-OC43-WT-Ribo-BGH-YCpBAC used for virus rescue. Numbered junctions detected by PCR (black arrows) and restriction sites used for TAR cloning are indicated. **(E)** PCR screening of assembled junctions 7/1, 1/2, 2/3, 3/4, 4/6, and 6/7 with water controls (dH2O) to the right of the panel. Primer pairs used for screening are listed in Table 2. Abbreviations: 100bp, 100 bp ladder; bp/kbp, base/kilobase pairs; *CmR*, chloramphenicol resistance gene; *poly(A)*, encoded A_34_ sequence; *URA3*, orotidine 5’-phosphate decarboxylase gene; WT, wild-type.

### Rescue of yeast assembled HCoV-OC43 wild-type and mClover reporter viruses

To recover infectious virus from these constructs, purified OC43-mClover or OC43-WT YCpBACs were co-transfected into 293T cells with a plasmid encoding *OC43-N* (Fig. 4A), as co-expression of N protein during the initial stages of coronavirus rescue supports virus rescue (15, 16, 61, 17). The transfected 293T cells were then co-cultured with more highly permissive BHK-21 cells (62, 63) to promote amplification of infectious virus (Fig. 4A). By the 6^th^ day in co-culture, the 293T cells had detached from the surface of the dish and the more adherent BHK-21 cells showed clear cytopathic effects (CPE) from infection with either OC43^YA^ or OC43-mClo^YA^ and mClover-positive nuclei were visible in >75% of the OC43-mClo^YA^ infected monolayer (Fig. 4A).

**Figure 4.**
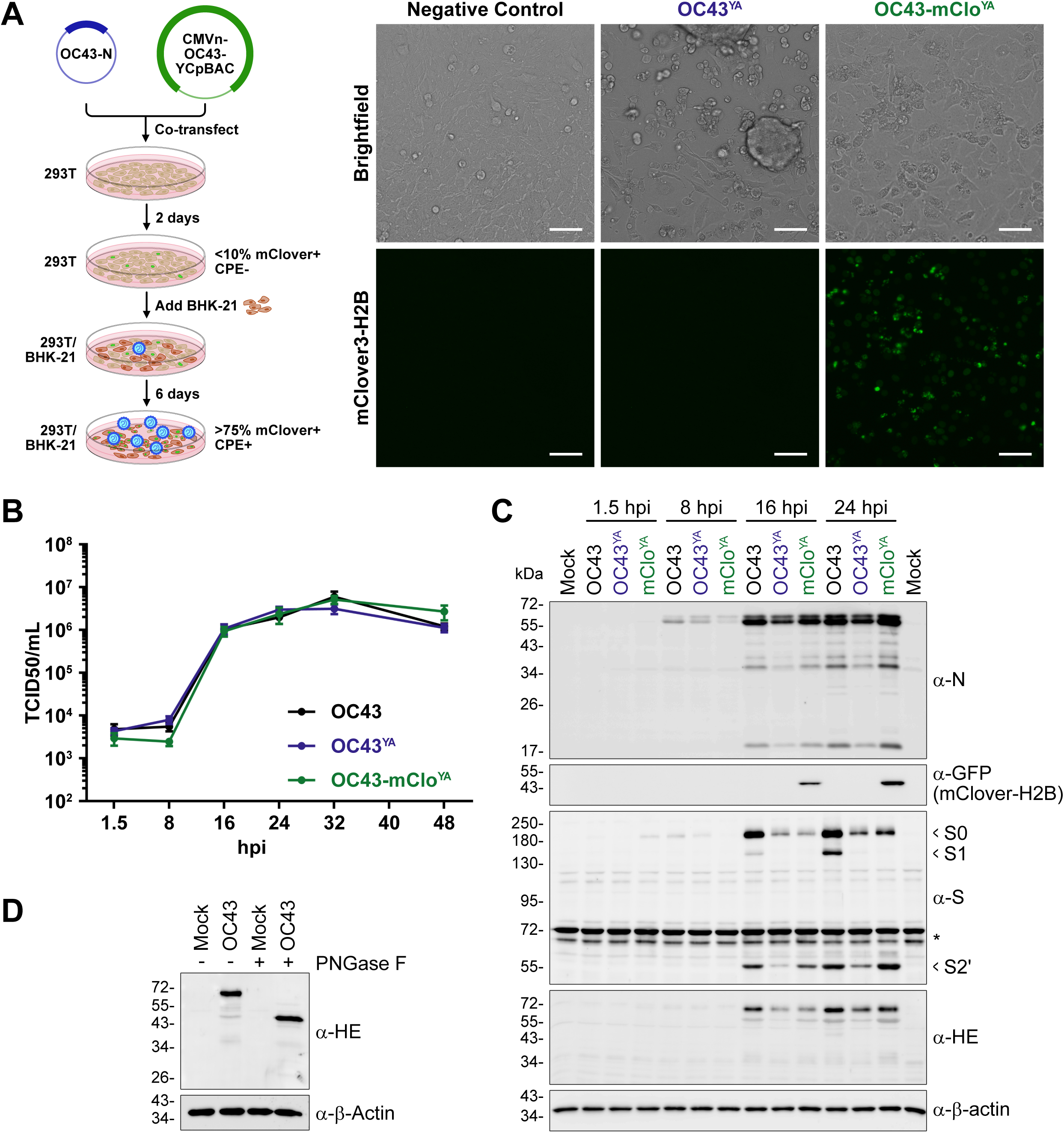
Yeast assembled HCoV-OC43 viruses replicate to comparable titers as wild-type HCoV-OC43 virus and produce similar amounts of viral proteins. **(A)** OC43-mClov^YA^ rescue procedure (left): Assembled CMVn-OC43-mClover plasmid and a plasmid encoding OC43-N were co-transfected into 293T (brown) cells for 2 days followed by re-seeding with BHK-21 (red) cells for 6 days to facilitate virus (blue) propagation. Live cell imaging (right) was performed on day 6 to examine the co-cultured cells for cytopathic effects (CPE; brightfield) and mClover3-H2B fluorescence (green nuclei). Scale bars = 50 µm. **(B)** Third passage OC43^YA^ and OC43-mClo^YA^ viruses were used to infect 293A cells at an MOI for 0.1 and the supernatants were collected at the indicated times and titered using BHK-21 cells. The data are plotted as the mean ± standard error of the mean from four independent replicates. **(C)** 293A cells were infected at an MOI of 0.1 with OC43 (ATCC; black), OC43^YA^ (yeast assembled OC43; blue), or OC43-mClo^YA^ (yeast assembled OC43-mClover; green) viruses for the indicated times. Protein lysates were subjected to SDS-PAGE and immunoblotted with the indicated antibodies where α-GFP antibodies were used to detect mClover3-H2B. Data presented are from one of four independent experiments. **(D)** Lysates from 293A cells infected with OC43 (24 hpi) were incubated with or without PNGase F prior to immunoblotting with antibodies against OC43-HE or β-Actin. Abbreviations: *, Non-specific proteins; GFP, green fluorescent protein; hpi, hours post-infection; PNGase F, peptide:N-glycosidase F; S0/S1, Spike subunit 0/1/2’; TCID50, 50% tissue culture infectious dose.

Compared to the ATCC strain of HCoV-OC43 (OC43), OC43-mClo^YA^ and OC43^YA^ replicated with similar kinetics over a 48 h time course of infection (Fig. 4B). We also compared viral protein production over the first 24 h of infection using HCoV-OC43-specific antibodies to N, S, and HE and a GFP antibody to detect mClover3-H2B expression (Fig. 4C). N protein accumulation (full-length,∼55 kDa) was observed as early as 8 hours post-infection (hpi) for all viruses and continued to increase over time with minimal differences in expression levels. This demonstrated that TRS* in the OC43-mClo^YA^ virus (Fig. 2D) did not significantly affect N protein production. Additionally, numerous smaller proteoforms of N (< 50 kDa) are produced from the yeast assembled viruses and similar to those observed during OC43(ATCC) infection, which is consistent with previous observations from infections with various coronaviruses (64–67). The OC43(ATCC) virus produced noticeably more HE and S than either of the two yeast assembled viruses. Multiple S isoforms were detected using the polyclonal S antibody, which are predicted to represent the S0, S1, and S2’ species based on their apparent molecular masses and likely arise from host protease processing of the S protein (Fig. 4C and (68)). We confirmed that the rabbit polyclonal HE antibody was specific for HE, as the protein was only detectable in infected cells and was sensitive to PNGase F treatment (Fig. 4D), consistent with previously reported *N*-glycosylation of HE (69, 70). We noted that despite being governed by identical TRS-B elements (compare TRS and TRS*, Fig. 2D), mClover3-H2B proteins were only detectable at 16 hpi by western blot, whereas N protein accumulated much earlier (Fig. 4C). Despite the observed differences in viral protein accumulation during viral replication, ultimately there was little impact on the amount of infectious virus released from cells (Fig. 4B).

### Insertion of an eighth sgRNA sequence alters coronavirus RNA transcriptional efficiencies

Next, we assessed viral genomic RNA (gRNA) and subgenomic RNA (sgRNA) production during infection with OC43^YA^ and OC43-mClo^YA^ over a 48 h time course (Fig. 5A). Reverse transcription was performed using random hexamers and oligo(dT) to amplify all sgRNA species ((-)sgRNA and sg-mRNA). The qPCR set used a common forward primer for all reactions that anneals in the 5’ leader sequence and unique reverse primers for each viral RNA with the resulting data normalized to *18S* rRNA and then plotted relative to *Orf1a* (gRNA). Overall, the trends in viral RNA production in cell infected with the three viruses were similar: The sgRNA transcripts proximal to *Orf1ab* (*NS2A*, *HE*, *S*, and *NS5A*) were at or below gRNA levels whereas the sgRNAs encoding *E*, *M*, and *N* were higher in abundance than gRNA (Fig. 5A). This is consistent with previously published RT-qPCR and transcriptomic analysis of HCoV-OC43 infection (71, 72). There were some differences in sgRNA accumulation at specific time points for both the OC43^YA^ and OC43-mClo^YA^ viruses compared to sgRNA accumulation from the OC43(ATCC) virus (Fig. 5A, indicated asterisks). The yeast assembled viruses produced higher levels of *E* (OC43^YA^), *M* (OC43^YA^), and *N* (OC43^YA^ and OC43-mClo^YA^) transcripts between 16-32 hpi when compared to the OC43(ATCC) infections (Fig. 5A, black asterisks). When comparing the two yeast assembled viruses, the OC43-mClo^YA^ virus produced less *E* and more *N* sgRNAs between 16-24 hpi and less *M* sgRNAs at 32 hpi (Fig. 5A, blue asterisks). Additionally, gRNA production (*Orf1a*, normalized to *18S* rRNA levels) was ∼6-fold higher in OC43-mClo^YA^ infected cells compared to those infected with OC43(ATCC) at 32 and 48 hpi (Fig. 5B, black asterisks) and slightly elevated in OC43-mClo^YA^ infected cells compared to OC43^YA^ gRNA levels in infected cells at 16 and 24 hpi (Fig. 5B, blue asterisks). The insertion of the reporter gene sequence and TRS* into the OC43-mClo^YA^ genome only moderately affected viral transcription (Fig. 5A). The *mClover3-H2B* sgRNA, which is produced from the authentic *N* TRS, only accumulated to levels equivalent to the *Orf1ab*-proximal genes *NS2A*, *HE*, *S*, and *NS5A*. This was unexpected considering the TRS sequence used (Fig. 2D) and location of the *mClover3-H2B* insertion near the 3’ end of the genome should allow for high sgRNA transcription (73, 74).

**Figure 5.**
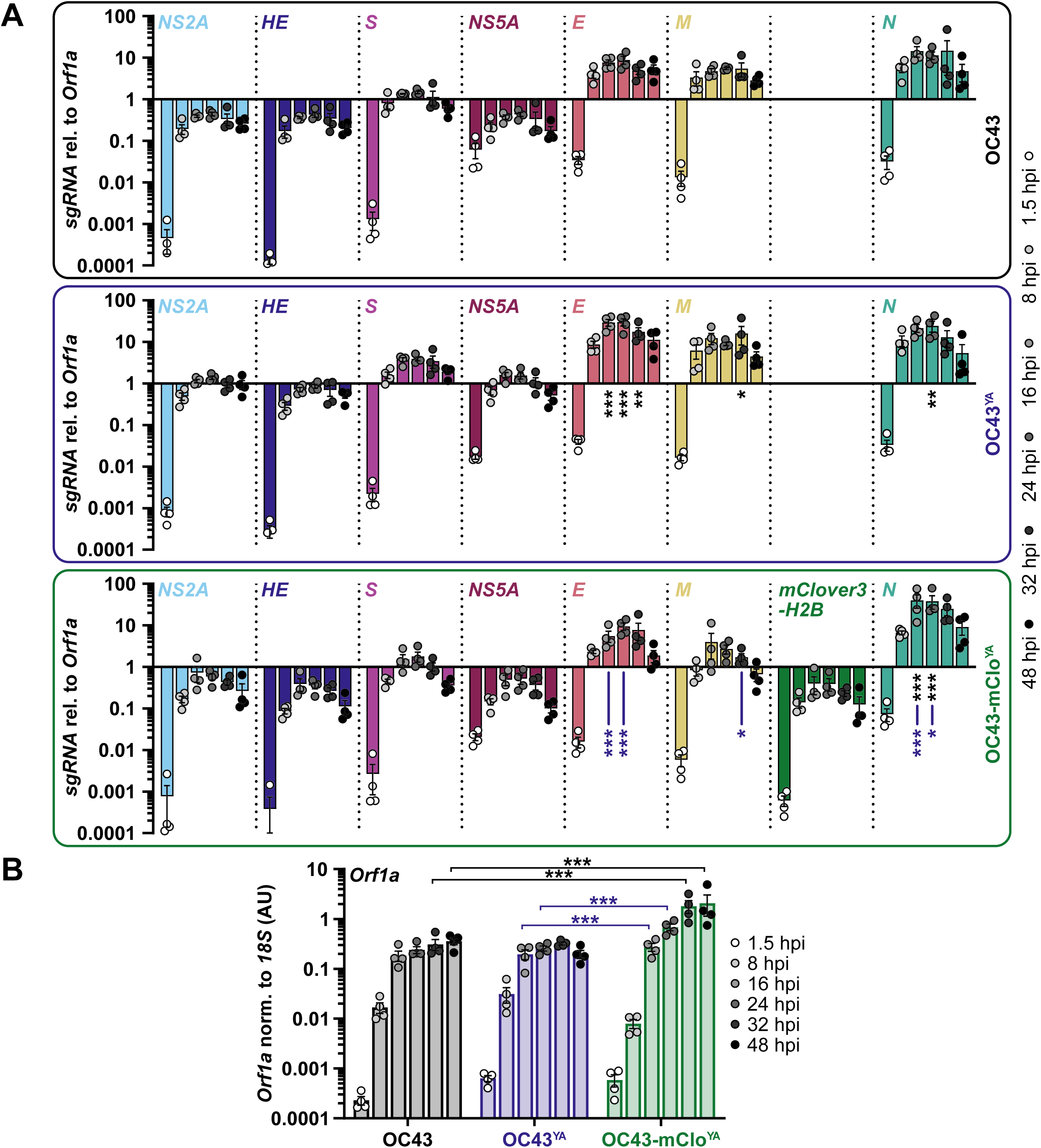
gRNA/sgRNA accumulation is altered by the insertion of the *mClover3-H2B* reporter gene. **(A)** 293A cells were infected at an MOI of 0.1 with OC43 (black box/top panel), OC43^YA^ (yeast assembled OC43; blue box/middle panel), or OC43-mClo^YA^ (yeast assembled OC43-mClover; green box/bottom panel) viruses for the indicated times. Total RNA was reverse transcribed and used for qPCR with a common forward primer that binds in the 5’ leader sequence and reverse primers that bind downstream of the leader (*Orf1a*) or downstream of ORF-specific transcription regulatory sequences. All data was normalized to *18S* rRNA and plotted relative to *Orf1a*. Data were plotted as the mean ± standard error of four independent experiments. Data points < 0.0001 are not shown. **(B)** *Orf1a* (gRNA) abundance normalized to *18S* rRNA and plotted as arbitrary units (AU). Same data set used in panel A and were plotted as the mean ± standard error of four independent experiments. Statistical comparisons were made between matched time points by 2-way ANOVA analysis. *p* values: * ≤ 0.05, ** ≤ 0.01, *** ≤ 0.001. Black asterisks represent differences compared to OC43 and blue asterisks represent differences between OC43^YA^ and OC43-mClo^YA^.

### OC43-mClo^YA^ allows for efficient and sensitive monitoring of viral replication

To more quantitatively measure viral replication kinetics, we used flow cytometry to measure N and mClover3-H2B protein expression in infected cells (Fig. 6). Cells from OC43(ATCC), OC43^YA^, or OC43-mClo^YA^ infections (MOI 0.1) were harvested at the same time points to those used for western blot analysis (Fig. 4C). These cells were fixed and stained with an antibody against OC43-N and fluorescence thresholds (gates) were set to yield <1% Alexa Fluor 647- or mClover3-H2B-positive mock infected cells. In all three infections, N expression was easily detectable by 8 hpi and began to plateau by 24 hpi (Fig. 6). The OC43-mClo^YA^ virus produced less N protein at 8 hpi and 16 hpi time points compared to the OC43(ATCC) virus but showed no N expression defect by 24 hpi. This flow cytometry approach was more sensitive compared to western blotting as we observed mClover3-H2B accumulation as early as 8 hpi, after which the fluorescence steadily increased over time. This allowed us to measure the accumulation of an N/mClover3-H2B double-positive population. In line with the RT-qPCR analysis (Fig. 5A), flow cytometry demonstrated that the mClover3-H2B protein has delayed expression kinetics relative to N and that the OC43-mClo^YA^ virus exhibited delayed accumulation of N relative to the OC43(ATCC) virus (Fig. 6B). However, by 24 hpi, N expression from the OC43-mClo^YA^ virus was equivalent to that of OC43(ATCC) and OC43^YA^ and endogenous mClover3-H2B expression more reliably identified infected cells (94% mClover+) compared to exogenous antibody staining (76% N+). This demonstrates that flow cytometric analysis of HCoV-OC43 infection is more sensitive than analysis by western blotting (compare Figs. 4C and 6B) and the OC43-mClo^YA^ reporter virus provides a more streamlined strategy to tracking infectivity in live cells or with only a fixation step needed prior to routine analysis.

**Figure 6.**
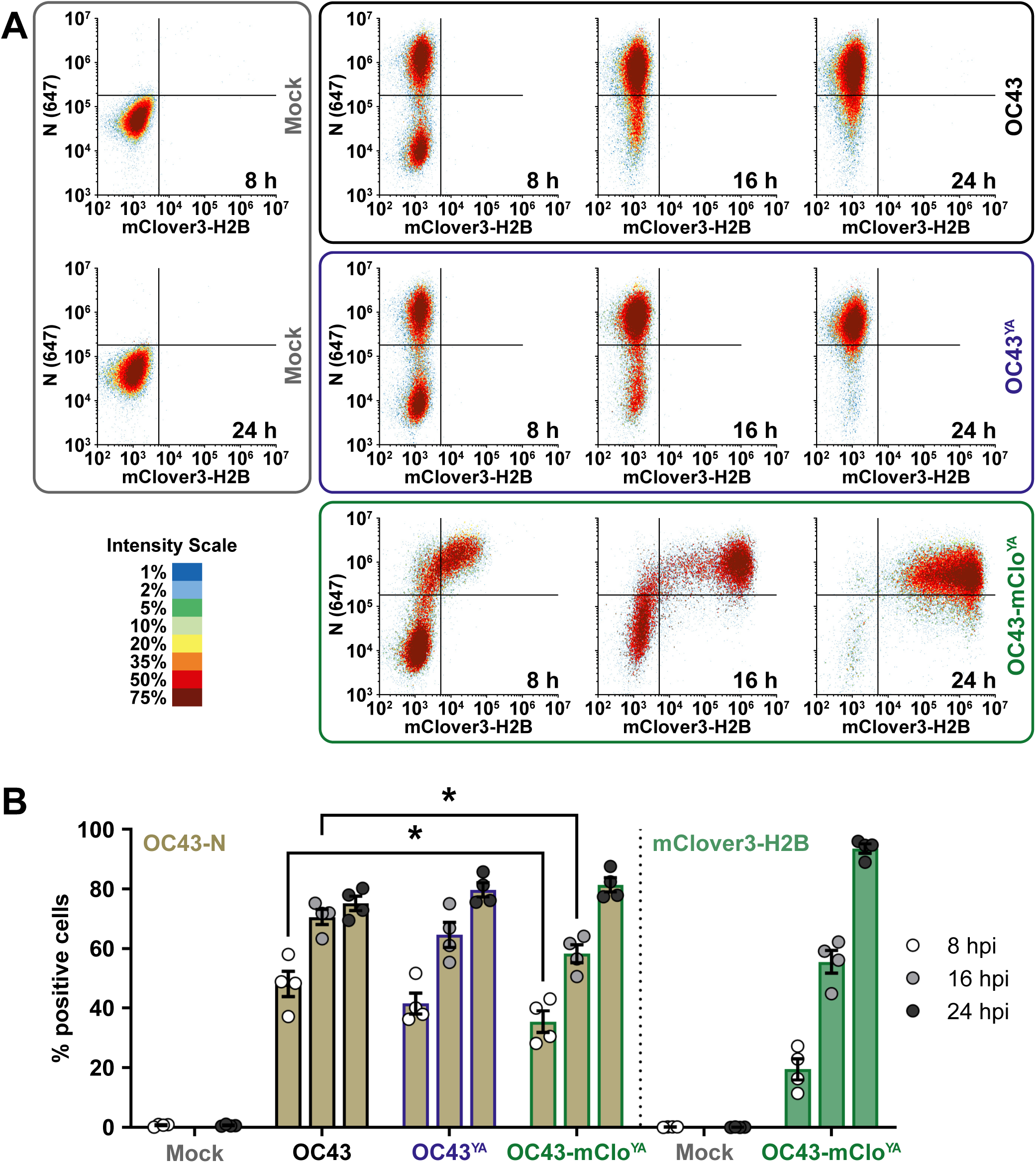
N protein accumulation is delayed during OC43-mClo^YA^ infection. (**A**) 293A cells were infected at an MOI of 0.1 with OC43 (black box/top panel), OC43^YA^ (yeast assembled OC43; blue box/middle panel), or OC43-mClo^YA^ (yeast assembled OC43-mClover; green box/bottom panel) viruses for 8, 16, or 24 hours post-infection (hpi) or mock infected for 8 or 24 hours (grey box/left panel). Cells were fixed, permeabilized, and stained with OC43-N antibodies prior to analysis by flow cytometry. Density plots of N (y-axis) vs. mClover3-H2B (x-axis) are shown from one representative experiment. **(B)** The data were analyzed for % positive single cells expressing OC43-N (light brown) or mClover3-H2B (light green) and plotted as the mean ± standard error of the mean from four independent experiments. Statistically significant differences in protein expression between different viruses are indicated (2-way ANOVA), * *p* ≤ 0.05. Abbreviations: 647, Alexa Fluor 647.

The reporter gene in the OC43-mClo^YA^ virus encodes a mClover variant which was engineered for enhanced photostability and brightness compared to EGFP (53). In addition to these beneficial properties, we also chose the mClover3-H2B fusion protein because it localizes exclusively to nuclei following expression (53). This is particularly advantageous when studying coronavirus biology by microscopy as the mClover3-H2B protein will not interfere with visualizing processes related to the cytoplasmic replication of coronaviruses. We used immunofluorescence microscopy to evaluate cytoplasmic replication compartment formation in cells infected with our WT strains or the OC43-mClo^YA^ virus. Using cells infected with an MOI of 0.1 for 16 h, we observed clear formation of replication compartments for all viruses based on dsRNA staining (Fig. 7). In cells infected with our OC43-mClo^YA^ virus, there were instances of dsRNA staining in cells that did not have detectable mClover3-H2B expression (Fig. 7, white arrowheads). This could be a consequence of the delayed mClover3-H2B expression kinetics compared to replication compartment formation. Future live cell imaging experiments are needed to determine if all dsRNA-positive cells eventually acquire mClover fluorescence.

**Figure 7.**
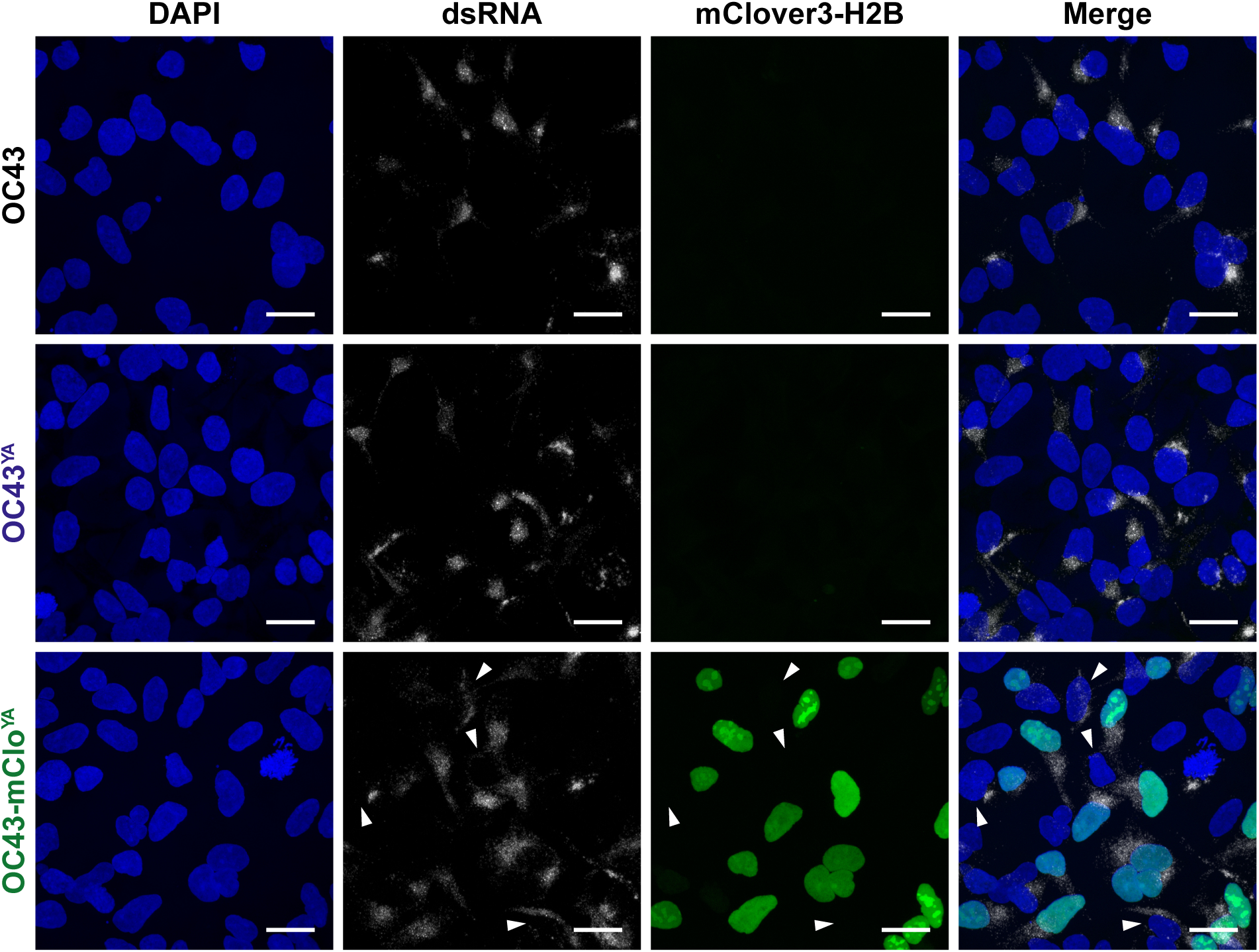
Replication compartment formation is unimpaired in cells infected with yeast assembled HCoV-OC43 viruses. 293A cells were infected at an MOI of 0.1 with OC43, OC43^YA^ (yeast assembled OC43), or OC43-mClo^YA^ (yeast assembled OC43-mClover) viruses for 16 hours prior to fixation, permeabilization, and staining with anti-dsRNA antibodies to stain replication compartments (white cytoplasmic puncta) and DAPI to stain cell nuclei (blue). mClover3-H2B is shown in green. White arrowheads denote dsRNA+/mClover-cells. All images were acquired using a confocal laser scanning microscope and are presented as maximum intensity projections from one of three independent experiments. Scale bars = 20 µm. Abbreviations: dsRNA, double-stranded RNA; H2B, histone H2B.

### Rapid evaluation of antiviral efficacy using OC43-mClo^YA^

Reporter viruses are useful tools for antiviral drug discovery. We developed a unique OC43-mClo^YA^ reporter virus, in part, to accelerate low-cost screening of novel compounds in a medium to high-throughput manner. To evaluate the OC43-mClo^YA^ virus as a screening tool for antiviral efficacy, we infected 293A, MRC-5, and BHK-21 cells at an MOI of 0.1 and treated these cells at 1 hpi with nirmatrelvir (PF-07321332), a potent coronavirus antiviral that targets Mpro (75). Matched cells and culture supernatants were used to monitor viral replication by flow cytometry and titering on BHK-21 cells, respectively. The OC43-mClo^YA^ virus infected and replicated in all cell lines tested (Fig. 8). Moreover, nirmatrelvir reduced viral titers by ∼3 log_10_ in all cell lines tested (Fig. 8A). These effects were mirrored by the flow cytometric analysis that showed clear reductions in the fluorescence intensity of infected cells following nirmatrelvir treatment (Figs. 8B-C). These data are consistent with the known antiviral activity of nirmatrelvir against HCoV-OC43 (75–77). The OC43-mClo^YA^ virus did not infect MRC-5 cells as readily as 293A or BHK-21 cells based on the intensity of mClover3-H2B staining in these cell populations (Figs. 8B-C), which limited the dynamic range for antiviral testing in MRC-5 cells with this reporter virus. Infection of the 293A and BHK-21 cells with OC43-mClo^YA^ produced far more and far brighter mClover+ cells compared to MRC-5 cells (Figs. 8B-C). Despite these differences in reporter expression, OC43-mClo^YA^ still replicated to high titers in the MRC-5 cells (Fig. 8A). These data together provide a proof-of-principle for the use of the OC43-mClo^YA^ virus for streamlined antiviral testing and highlight the utility of synthetic biology techniques for the generation and study of mutant coronaviruses.

**Figure 8.**
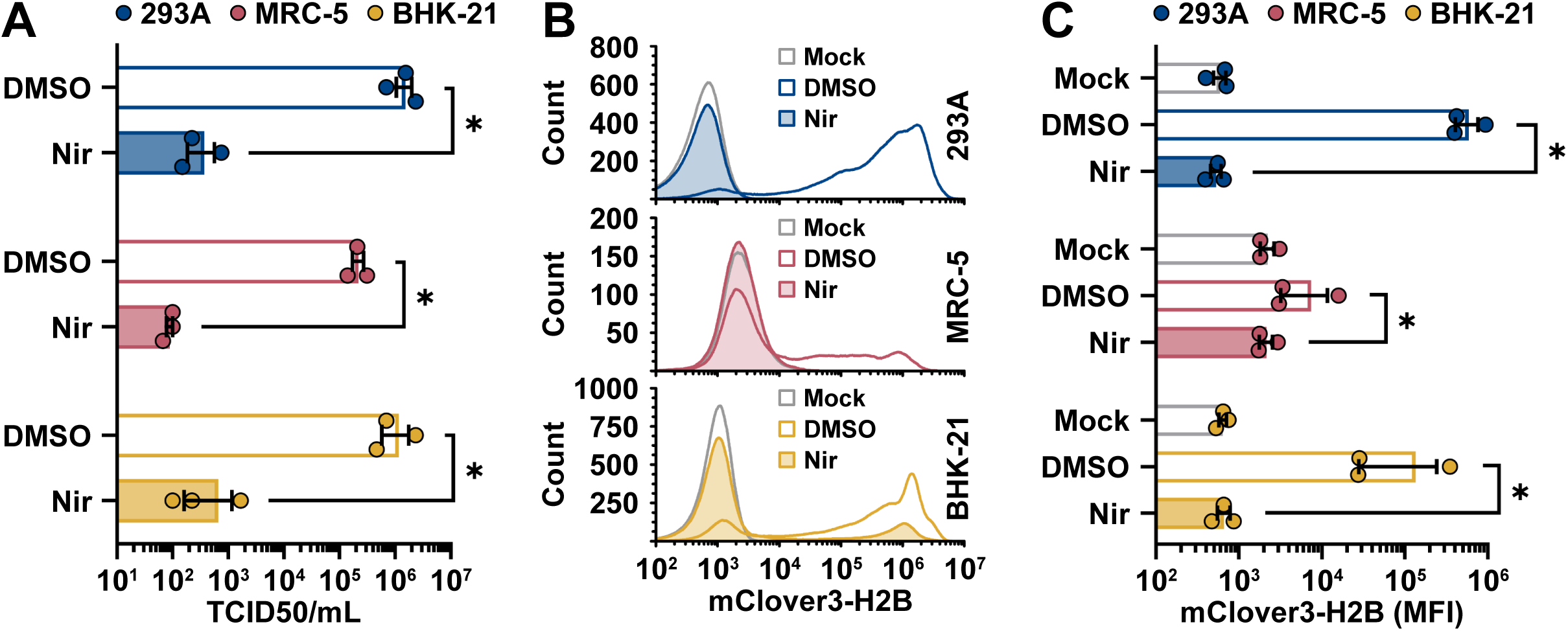
OC43-mClo^YA^ is responsive to antiviral treatment. 293A, MRC-5, or BHK-21 cells were mock infected or infected at an MOI of 0.1 with OC43-mClo^YA^ (yeast assembled OC43-mClover) for 1 hour prior to a medium change to 2.5% DMEM+ Pen/Strep/Gln supplemented with 0.1% DMSO or 1 µM nirmatrelvir (Nir). At 24 hours post-infection, the supernatants were collected for titering on BHK-21 cells (**A**). The cell monolayers were collected and fixed prior to analysis of mClover3-H2B expression by flow cytometry (**B**) with the average mClover3-H2B median fluorescence intensity (MFI) from mock or infected cells plotted in panel **C**. Data were plotted as the mean ± standard error of three independent experiments. Statistical comparisons were made by 2-way ANOVA analysis. *P* values: * ≤ 0.05. Abbreviations: DMSO, dimethyl sulfoxide; TCID50, 50% tissue culture infectious dose.

## DISCUSSION

Once established, reverse genetics systems provide well-defined clonal genetic material that can be readily propagated and mutagenized for fundamental studies of viral biology and a variety of biomedical applications. Here, we report the creation of an updated reverse genetics system for the endemic, seasonal, betacoronavirus HCoV-OC43. We used TAR methodology to capture fragments of the ∼31 Kb (+)ssRNA HCoV-OC43 genome to store as sequence-validated dsDNA parts, and performed combinatorial assembly in yeast, yielding an intact dsDNA copy of the HCoV-OC43 genome that could initiate a full viral gene expression program upon introduction into cell lines (Fig. 4). Moreover, we provide proof-of-concept for the engineering of HCoV-OC43 reporter viruses encoding additional sgRNAs without altering the coding regions of neighboring coronavirus genes. We thoroughly evaluated the yeast assembled, recombinant HCoV-OC43 viruses, demonstrating comparable viral gene expression, susceptibility to antivirals, and fitness in cell culture compared to its natural progenitor (Figs. 4-8). Thus, our approach to coronavirus genome assembly provides a viable path to generate recombinant HCoV-OC43 viruses.

Coronaviruses rely on discontinuous transcription to create (-)sgRNAs that are converted into sg-mRNAs that encode structural proteins and accessory proteins. This process relies on sequence complementarity between nascent (-)sgRNA TRS-B hairpin sequences and parental (+)gRNA TRS-L hairpin sequences, which enables template switching by the viral RdRp. Despite evidence that the RdRp favours generation of short (-)sgRNA by template switching at the first TRS-B that it encounters, the precise rules that govern template-switching preferences and the ultimate stoichiometry of sgRNA products remain poorly understood. Considering the similarity of TRS-B sites across the HCoV-OC43 genome, we reasoned that insertion of a reporter gene linked to its own TRS-B should enable efficient production of an additional sg-mRNA encoding the reporter gene (Fig. 2D). Our study demonstrated the feasibility of this approach for HCoV-OC43, whereby linking the reporter gene in the OC43-mClo^YA^ virus to the *N* TRS-B resulted in efficient mClover3-H2B expression (Fig. 6); with minimal effects on N expression (Figs. 4C and 6B) from the *N* sg-mRNA regulated by the 23 base TRS* (Fig. 2D). The OC43-mClo^YA^ virus displayed equivalent replication kinetics and overall fitness in cell culture compared to the OC43(ATCC) virus (Fig. 4B). Our observations were consistent to those made using the MHV system where the TRS/luciferase gene insertions were well-tolerated and recombinant MHVs replicated similarly to wild-type virus with modest attenuation of upstream sgRNA expression (10). This prior work also demonstrated that the genomic position and sequence of the inserted gene can influence reporter expression and stability (10), making it necessary to empirically evaluate future heterologous gene insertions into the HCoV-OC43 genome.

The sgRNAs encoding *mClover3-H2B* accumulated to lower levels than the *E* and *M* sgRNAs, as well as the *N* sgRNAs containing identical core TRS-B sites (Fig. 5). We speculate that this difference in sgRNA levels could stem from less efficient template switching at the *mClover3-H2B* TRS-B compared to the other authentic TRS-Bs, perhaps because of RNA sequences between the TRS and *mClover3-H2B* start codon as a result of our cloning strategy. Interestingly, this conclusion differs from those of previous studies using mouse hepatitis virus defective interfering (DI) RNA constructs that demonstrate that inserted sequences of up to ∼1.5 kb upstream or downstream of a DI RNA TRS had little impact on sgRNA production (78). This suggests there are differences in how the RdRp interacts with TRS-flanking sequences in the context of the full-length gRNA compared to artificial DI RNA constructs. Coronavirus sgRNA abundance is inversely proportional to the length and the number of internal TRSs present within the transcript (73, 74). We did not observe a stepwise reduction in upstream sgRNA/gRNA abundance during infection with the OC43-mClo^YA^ virus that one might expect from the insertion of an additional TRS-B; conversely gRNA levels were increased in OC43-mClo^YA^ infected cells compared to WT virus infection (Fig. 5B). The unexpected regulation of *mClover3-H2B* sgRNA transcripts would suggest that there are more complex regulatory processes at play during authentic viral infection. Future work will alter the RNA sequence present in the OC43-mClo^YA^ genome to attempt to enhance reporter gene expression while also providing insight into how TRS context affects coronavirus transcription.

A benefit of the TAR methodology is the flexibility of altering viral sequences through fragment-mediated assembly. This allows for the rapid generation of recombinant viruses from clinical isolates as demonstrated with SARS-CoV-2 and HCMV (45, 48). Not only could this allow for the study of emerging viruses with reverse genetics, but it may also allow for swapping of sequences to study gene variants or strain-specific differences. HCoV-OC43, as well as other human coronaviruses, circulate in a recurrent, seasonal pattern due to short-lived host immunity (79, 80). This can be attributed, in part, to adaptive evolutionary changes observed in the HCoV-OC43 Spike protein sequence over time (81); where the presence of HCoV-OC43 antibodies can reduce the severity of subsequent coronavirus infection (82, 83). The TAR methodology allows for the generation of pools of HCoV-OC43 viruses based on naturally occurring strains to determine which strains elicit superior cross-neutralizing antibody responses in animal models for possible vaccine development. Likewise, naturally occurring sequences could be engineered into HCoV-OC43 genomes via TAR to test for resistance to current or novel direct-acting antivirals.

## MATERIALS AND METHODS

### Mammalian cells and viruses

Human embroyonic kidney 293A, human embroyonic kidney 293T, and human lung fibroblast MRC-5 cells were grown in Dulbecco’s modified Eagle’s medium (DMEM; Thermo Fisher, 11965118) supplemented with heat-inactivated 10% fetal bovine serum (FBS, Thermo Fisher, A31607-01), 100 U/mL penicillin, 100 µg/mL streptomycin, and 2 mM L-glutamine (Pen/Strep/Gln; Thermo Fisher, 15140122 and 25030081). Baby hamster kidney (BHK-21) cells were maintained as above yet supplemented with 5% FBS. All cells were maintained at 37°C in 5% CO_2_ atmosphere.

Stocks of human coronavirus OC43 (HCoV-OC43; ATCC, VR-1558) and recombinant HCoV-OC43 were propagated in BHK-21 cells. Cells were infected at a MOI of 0.05 for 1 h at 37°C in serum-free DMEM. After 1 h, the infected cells were maintained in DMEM supplemented with 1% FBS and Pen/Strep/Gln until cytopathic effect was complete. Upon harvest, the culture supernatant was centrifuged at 1000 x g for 5 min at 4°C, aliquoted, and stored at −80°C. Viral titers were measured using median tissue culture infectious dose (TCID50) assays. Following the five-fold serial dilutions of the samples, BHK-21 cells were infected for 1 h at 37°C, followed by replacement of inocula with 1% FBS in DMEM+Pen/Strep/Gln. Viral titers were calculated using the Spearman-Kärber method (84).

### Plasmids

The plasmid pCC1-BACYCp-URA3 (YCpBAC; (57)) was used for all transformation-associated recombination (TAR) cloning. The plasmids pKanCMV-mClover3-10aa-H2B (mClover3 fused to a C-terminal histone H2B; Addgene, Plasmid#74257 (53)), pKanCMV-mRuby3-10aa-H2B (mRuby3 fused to a C-terminal histone H2B; Addgene, Plasmid#74258 (53)), mCardinal-C1 (Addgene, Plasmid#54799 (85)), EBFP2-C1 (Addgene, Plasmid#54665), and FLuc Control Template (NEB) were used as templates for amplification of the respective reporter ORFs. The CMV promoter (CMVpro) and BGH poly(A) signal (BGH) sequences were obtained from pcDNA3.1(+). The TAR cloning plasmids pUC57-N-TRS and pUC57-HDV-T7term encoding the *OC43-N* TRS or hepatitis delta virus (HDV) ribozyme and T7 terminator sequences, respectively, were synthesized by GenScript. A codon-optimized HCoV-OC43 N expression plasmid, pTwist-OC43-N(CO) (Addgene, Plasmid#151960), was co-transfected with assembled HCoV-OC43 plasmids during viral rescue with 293T cells. All plasmids generated for this study are included in Table 3.

**Table 3:**
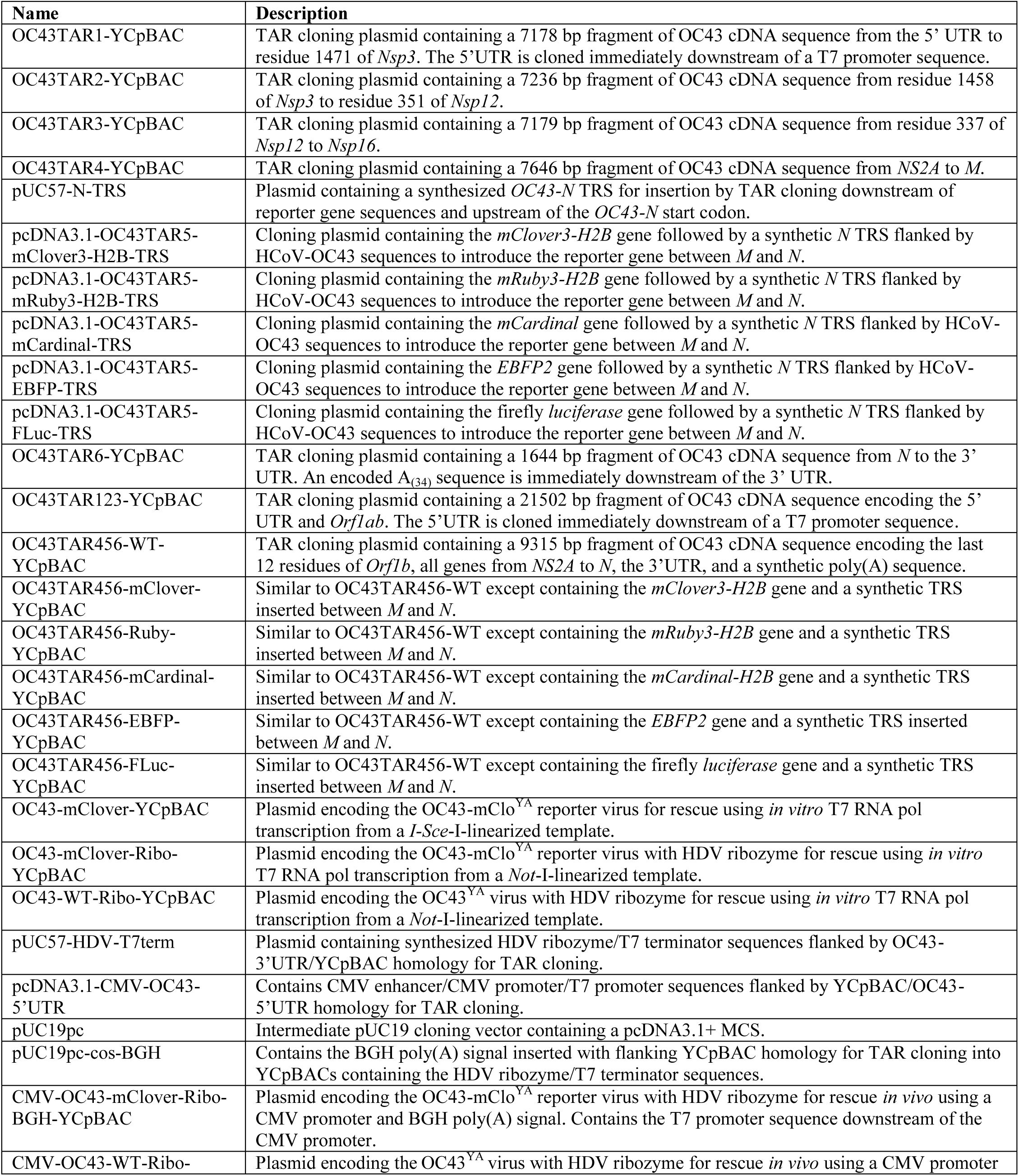
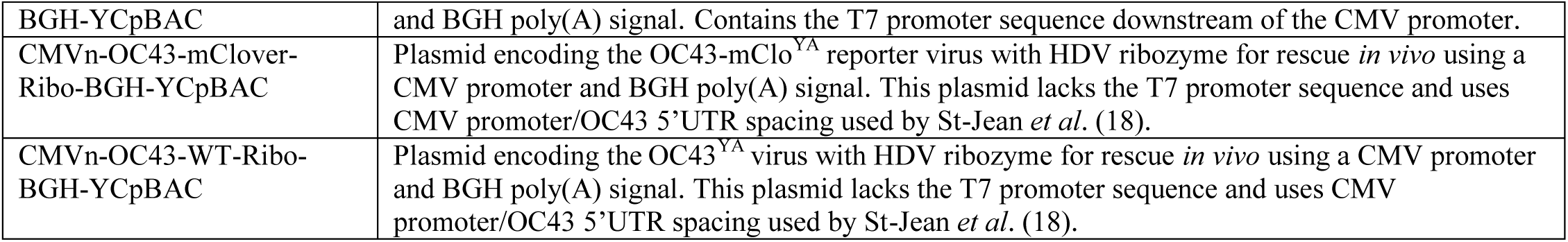
Plasmids generated for HCoV-OC43 TAR assembly.

### Yeast and bacterial cell propagation and transformation

*Saccharomyces cerevisiae* strain VL6-48N (*MATα*, *his3-Δ200*, *trp1-Δ1*, *ura3-Δ1*, *lys2*, *ade2-101*, *met14*, *psi*+*cir^0^*) (86) was used for all yeast transformation experiments following previously established protocols (87, 88) with the following changes: Yeast cells were grown in yeast extract peptone dextrose (YEPD) medium to a density of 2-4 x 10^7^ cells/mL, the working concentration of zymolyase (80 µg/mL; MP Biomedicals, 08320921) was empirically determined to yield near complete spheroplast generation after 30-40 min as determined by visualization of spheroplasts in 1 M sorbitol compared to those in 2% SDS using a hemocytometer, each 200 µL spherolplast suspension contained: < 50 ng amplified pCC1BACYCp-URA (YCpBAC) plasmid (linear capture plasmids), < 3 µg linear viral genomic cDNA amplicons/restriction fragments and/or recombinant DNA, and 5-10 µg of salmon sperm DNA (Thermo Fisher, 15632011), and incubations in PEG 8000 solution or SOS solution were 20 min and 30 min, respectively. Transformed spheroplasts resuspended in melted SORB-TOP-URA and plated onto SORB-URA plates prior to incubation at either room temperature or 30°C until colonies formed (87).

*Escherichia coli* strain Stbl3 (Thermo Fisher, C7373) were used to maintain and propagate all plasmids in bacteria. An overnight culture of these cells was rendered electrocompetent by growing to log-phase followed by three washes and resuspension in ice-cold 10% glycerol. To transfer YCpBACs into Stlb3 *E. coli*, total yeast DNA obtained from cells following overnight liquid culture in SC-URA medium was purified using the Gentra/Puregene Yeast/Bacteria Kit (QIAGEN, 158567). Yeast DNA was electroporated into Stbl3 cells using a Gene Pulser II electroporation system (Bio-Rad) set to 2.5 kV/25 µF/200 Ω with 0.2 cm electroporation cuvettes (VWR, 89047-208).

### Synthesis and capture of HCoV-OC43 genome fragments

To generate YCp plasmids, the HCoV-OC43 genome was divided into five TAR fragments (1, 2, 3, 4, and 6). Corresponding primer pairs (OC43-TARxF/OC43-TARxR, Table 1) for each TAR fragment were used to first amplify the YCpBAC plasmid to generate linear capture plasmids with 45 bp of sequence homology associated with the 5’ and 3’ ends of each viral genomic fragment using KOD Xtreme Hot Start DNA Polymerase (KODpol; Sigma-Aldrich, 71975-M) following the manufacturer’s instructions for two-step cycling. These primers were designed to incorporate either *Not*I (TAR1 and TAR7) or *I-Sce*I (TAR2-6) restriction sites between the YCpBAC and HCoV-OC43 sequences to facilitate the release of HCoV-OC43 sequences for TAR cloning.

Viral RNA from HCoV-OC43 virions (ZeptoMetrix, 0810024CF; ATCC, VR-1558) was isolated using the RNeasy Plus Mini Kit (QIAGEN) and reverse transcribed into cDNA fragments using gene-specific primers 1/2R, 2/3R, 3/4R, 5/6R (Table 1) or oligo(dT)_20_ and the SuperScript III First-Strand Synthesis System (Invitrogen, 18080-051) with a modified protocol for long cDNA synthesis (89). cDNAs were amplified using PCR with primers TAR1ampF/1/2R (TAR1), 1/2F/2/3R (TAR2), 2/3F/3/4R (TAR3), 3/4F/5/6R (TAR4), or 4/5F/TAR6ampR (TAR6) (Table 1) to generate dsDNA fragments of the HCoV-OC43 genome using KODpol following the manufacturer’s instructions.

To generate YCpBACs containing single TAR fragments, linear capture plasmids were co-transformed into yeast with cDNA amplicons spanning the desired region of the viral genome to generate the plasmids: OC43TAR1-YCpBAC, OC43TAR2-YCpBAC, OC43TAR3-YCpBAC, OC43TAR4-YCpBAC, and OC43TAR6-YCpBAC.

The TAR5(mClover3-H2B) dsDNA fragment was generated by PCR with KODpol using the OC43-TAR5F-mClover/OC43-TAR5R-mClover primers using the pKanCMV-mClover3-10aa-H2B plasmid as a template. This amplicon was digested with *Hind*III and phosphorylated with T4 polynucleotide kinase (New England Biolabs (NEB), M0201S) and cloned into pcDNA3.1+ digested with *Eco*RV and *Hind*III restriction sites to create pcDNA3.1-TAR5-mClover3-H2B. Since the positioning of the TAR5 reporter gene will be downstream of the authentic *N* transcription regulatory sequence (TRS), a second copy of the *N* TRS (TRS*, Fig. 2D) was inserted downstream of the reporter genes to produce the sgRNAs encoding *N* in reporter viruses. A *Bam*HI/*Afl*II restriction fragment containing the duplicated TRS from pUC57-N-TRS was ligated into *Bam*HI/*Afl*II digested pcDNA3.1-TAR5-mClover3-H2B to generate pcDNA3.1-TAR5-mClover3-H2B-TRS. The remaining TAR5 reporter fragments were generated by *Age*I/*Bam*HI digestion of pKanCMV-mRuby3-H2B, mCardinal-C1, EBFP2-C1, or a PCR amplicon of FLuc Control Template (AgeI-TAR5F-FLuc/BamHI-TAR5R-FLuc, KODpol) and ligated into pcDNA3.1-OC43TAR5-mClover3-H2B-TRS cut with the same enzymes to generate: pcDNA3.1-OC43TAR5-mRuby3-H2B-TRS, pcDNA3.1-OC43TAR5-mCardinal-TRS, pcDNA3.1-OC43TAR5-EBFP2-TRS, and pcDNA3.1-OC43TAR5-FLuc-TRS. The wild-type (WT)/reporter-less TAR fragment was generated by PCR of viral cDNA using KODpol and the primers OC43-TAR5F-WT/OC43-TAR5R-WT.

### Screening of yeast assemblies by colony PCR

Yeast colonies from the SORB-URA transformation plates were patched onto SC-URA plates and incubated overnight at 30°C. The patched yeast were lysed in 20 µL 0.05% SDS at 95°C for 15 min or 0.25 mg/mL zymolyase in 50 mM Tris-Cl (pH 7.5)/25% glycerol at 37°C for 15 min. 1-5 µL of lysate was screened using 25 µL *Taq* DNA Polymerase with ThermoPol Buffer (NEB, M0267L) reactions following the manufacturer’s protocol with 500 nM primers (Table 2). PCRs were analyzed following DNA gel electrophoresis using 1.8% agarose/TAE gels with a Wide Mini-Sub Cell GT apparatus and 26-well combs (Bio-Rad, 1704469 and 1704457).

### Assembly of HCoV-OC43 sequences in yeast

Partial assemblies of OC43TAR123-YCpBAC (containing *Orf1ab*) or OC43TAR456— Ruby3-H2B were assembled in yeast using equimolar ratios of the individual TAR plasmids and 5-fold less YCpBAC capture plasmid. The individual TAR fragments were linearized for assembly following restriction digestion or PCR amplification using KODpol. A truncated YCpBAC capture plasmid generated by KODpol PCR with OC43-TAR7F-trunc/OC43-TAR7R-trunc primers was used for the partial assemblies. Partial assemblies of OC43TAR456-WT, mClover, mCardinal, EBFP, or FLuc YCpBACs were performed using a *Bsi*WI-digested OC43TAR456-mRuby-H2B-YCpBAC (Fig. 2B) co-transformed with *I-Sce*I-digested pcDNA3.1-OC43TAR5 reporter plasmids. Full assemblies (Fig. 3A) were performed by co-transforming equimolar ratios of *Sgr*AI/*I-Sce*-I-digested OC43TAR123-YCpBAC and *Srf*I/*Sgr*AI-digested OC43TAR456-mClover-YCpBAC (Fig. 2A-B) with 10-fold less truncated YCpBAC to generate OC43-mClover-YCpBAC.

Yeast colonies were screened for YCpBACs containing the desired DNA junctions using *Taq* PCRs as indicated above. Completed assemblies were verified by restriction analysis, Sanger sequencing (Genewiz), and Oxford Nanopore sequencing (Plasmidsaurus).

### Insertion of transcriptional control and RNA modification elements using TAR cloning

Following assembly of the full-length OC43-mClover-YCpBAC, sequences encoding the HDV ribozyme and T7 terminator were inserted immediately 3’ of the encoded poly(A) sequence (Fig. 3A). The HDV ribozyme/T7 terminator sequences were excised from pUC57-HDV-T7term using *Esp*3I and co-transformed with the OC43-mClover-YCpBAC digested with *I-Sce*I into yeast for TAR assembly to generate OC43-mClover-Ribo-YCpBAC. *Bam*HI/*I-Sce*-I-digested OC43-mClover-YCpBAC was co-transformed with a PCR amplicons containing WT HCoV-OC43 sequence spanning the *M*-*N* genes (4/5F/5/6R, OC43 cDNA, KODpol) and the sequences encoding the HDV ribozyme/T7 terminator (6/7F/6/7R, OC43-mClover-Ribo-YCpBAC) to generate OC43-WT-Ribo-YCpBAC (Fig. 3A).

To modify the YCpBACs for mammalian cell rescue, CMVpro and BGH sequences were inserted into OC43-mClover-Ribo-YCpBAC (Fig. 3A). CMVpro: A *Xho*I/*Bgl*II fragment from OC43TAR1-YCpBAC containing the OC43 5’UTR was cloned into pcDNA3.1+ digested with the same enzymes. The resulting plasmid was digested with *Srf*I and a blunted (NEB Quick Blunting Kit) *Mlu*I/*Sac*I digest of pcDNA3.1+ (containing the CMVpro sequence) ligated into the backbone to create pcDNA3.1-CMV-OC43-5’UTR. BGH: pUC19 digested with *Nde*I/*Xba*I was ligated with a *Nde*I/*Xba*I insert containing the pcDNA3.1+ MCS to generate pUC19pc. The *Eco*RI/*Hpa*I *cos*-containing fragment from YCpBAC was ligated into pUC19pc digested with *Eco*RI/*Pvu*II to create pUC19pc-cos. A PCR amplicon of the pcDNA3.1+ BGH (BD206/BD207, containing BGH, KODpol) was phosphorylated and blunt-end ligated into pUC19pc-cos digested with *Not*I and blunted to generate pUC19pc-cos-BGH. The OC43-mClover-Ribo-YCpBAC was digested with *Not*I and co-transformed into yeast with the *Xho*I/*Bgl*II fragment of pcDNA3.1-CMV-OC43-5’UTR and the *Sfo*I/*Sal*I fragment of pUC19pc-cos-BGH to generate CMV-OC43-mClover-Ribo-BGH-YCpBAC. To remove the T7 promoter sequence in CMV-OC43-mClover-Ribo-BGH-YCpBAC and shorten the spacing between CMVpro and the OC43 5’UTR (18), a pair of oligonucleotides (CMVn-OC43F/CMVn-OC43R) were annealed and filled-in with DNA Polymerase I, Large Fragment (Invitrogen, 18012021) and co-transformed with *Not*I-digested CMV-OC43-mClover-Ribo-BGH-YCpBAC to generate CMVn-OC43-mClover-Ribo-BGH-YCpBAC (Fig. 3A).

The WT version of this plasmid was created by co-transforming *Bam*HI-digested CMV-OC43-mClover-Ribo-BGH-YCpBAC with a PCR amplicon of WT HCoV-OC43 sequence (4/5F/6/7R, OC43-WT-Ribo-YCpBAC, KODpol) into yeast to generate CMV-OC43-WT-Ribo-BGH-YCpBAC (Fig. 3A). This plasmid was then digested with *Not*I and co-transformed with annealed and filled-in CMVn-OC43F/CMVn-OC43R to generate CMVn-OC43-WT-Ribo-BGH-YCpBAC (Fig. 3A).

Yeast colonies were screened for YCpBACs containing the desired DNA junctions using *Taq* PCRs as indicated above. Completed assemblies were verified by restriction analysis, Sanger sequencing (Genewiz), and Oxford Nanopore whole plasmid sequencing (Plasmidsaurus). Single-nucleotide polymorphisms in CMVn-OC43-WT-Ribo-BGH-YCpBAC (WT) and CMVn-OC43-mClover-Ribo-BGH-YCpBAC (mClover) plasmids compared to a reference strain of HCoV-OC43 (GenBank accession ON376724) are listed in Table 4.

**Table 4:**
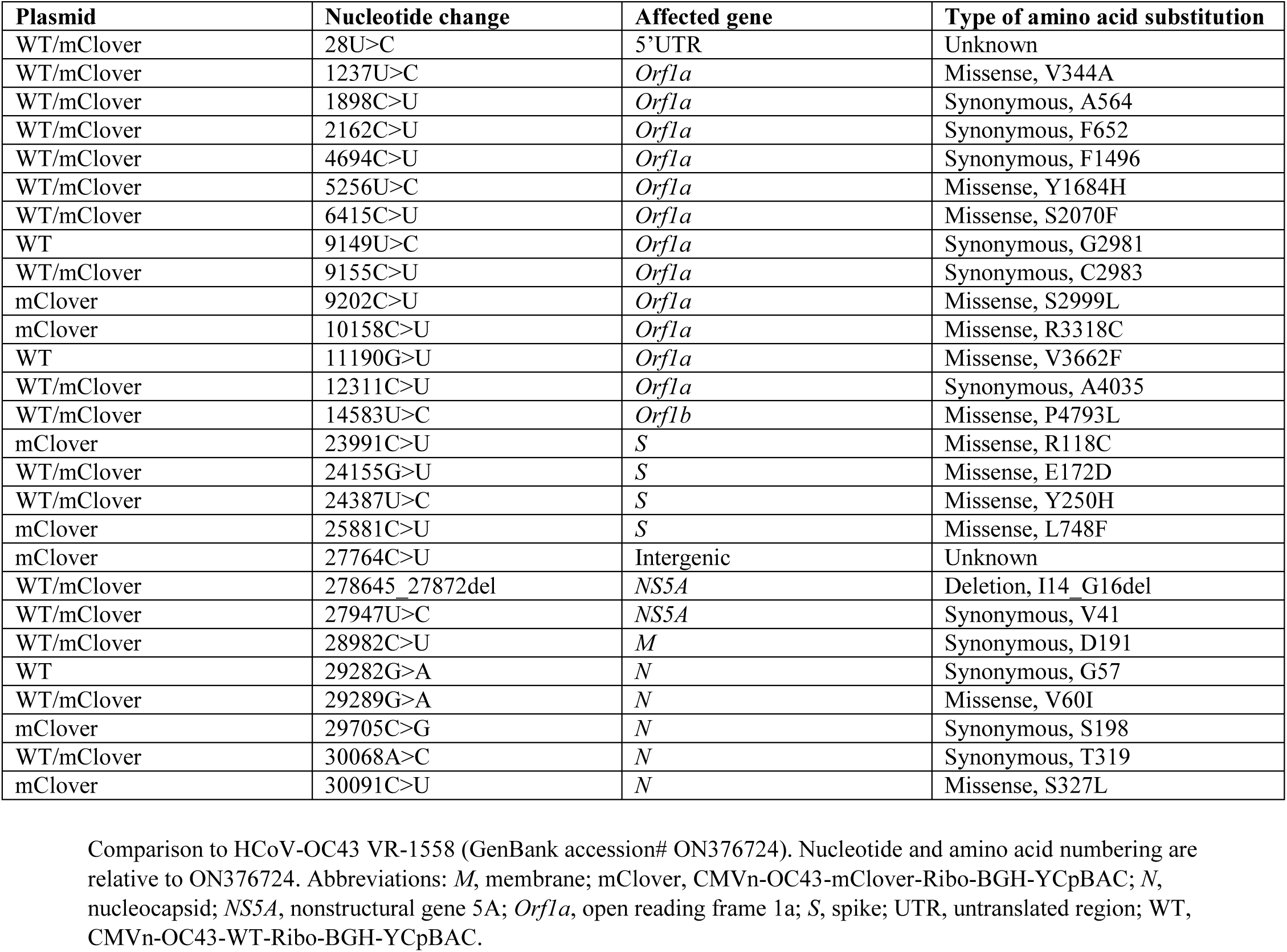
Single-nucleotide polymorphisms in YCpBACs used for virus rescue compared to a reference stain of HCoV-OC43.

| Plasmid | Nucleotide change | Affected gene | Type of amino acid substitution |
| --- | --- | --- | --- |
| WT/mClover | 28U>C | 5'UTR | Unknown |
| WT/mClover | 1237U>C | *Orf1a* | Missense, V344A |
| WT/mClover | 1898C>U | *Orf1a* | Synonymous, A564 |
| WT/mClover | 2162C>U | *Orf1a* | Synonymous, F652 |
| WT/mClover | 4694C>U | *Orf1a* | Synonymous, F1496 |
| WT/mClover | 5256U>C | *Orf1a* | Missense, Y1684H |
| WT/mClover | 6415C>U | *Orf1a* | Missense, S2070F |
| WT | 9149U>C | *Orf1a* | Synonymous, G2981 |
| WT/mClover | 9155C>U | *Orf1a* | Synonymous, C2983 |
| mClover | 9202C>U | *Orf1a* | Missense, S2999L |
| mClover | 10158C>U | *Orf1a* | Missense, R3318C |
| WT | 11190G>U | *Orf1a* | Missense, V3662F |
| WT/mClover | 12311C>U | *Orf1a* | Synonymous, A4035 |
| WT/mClover | 14583U>C | *Orf1b* | Missense, P4793L |
| mClover | 23991C>U | *S* | Missense, R118C |
| WT/mClover | 24155G>U | *S* | Missense, E172D |
| WT/mClover | 24387U>C | *S* | Missense, Y250H |
| mClover | 25881C>U | *S* | Missense, L748F |
| mClover | 27764C>U | Intergenic | Unknown |
| WT/mClover | 278645_27872del | *NS5A* | Deletion, I14_G16del |
| WT/mClover | 27947U>C | *NS5A* | Synonymous, V41 |
| WT/mClover | 28982C>U | *M* | Synonymous, D191 |
| WT | 29282G>A | *N* | Synonymous, G57 |
| WT/mClover | 29289G>A | *N* | Missense, V60I |
| mClover | 29705C>G | *N* | Synonymous, S198 |
| WT/mClover | 30068A>C | *N* | Synonymous, T319 |
| mClover | 30091C>U | *N* | Missense, S327L |

Comparison to HCoV-OC43 VR-1558 (GenBank accession# ON376724). Nucleotide and amino acid numbering are relative to ON376724. Abbreviations: *M*, membrane; mClover, CMVn-OC43-mClover-Ribo-BGH-YCpBAC; *N*, nucleocapsid; *NS5A*, nonstructural gene 5A; *Orf1a*, open reading frame 1a; *S*, spike; UTR, untranslated region; WT, CMVn-OC43-WT-Ribo-BGH-YCpBAC.

### Rescue of infectious yeast assembled HCoV-OC43 recombinant viruses

6-wells of 293T cells seeded at ∼500,000 cells per well were co-transfected with 1 µg of CMVn-OC43-mClover-Ribo-BGH-YCpBAC or CMVn-OC43-WT-Ribo-BGH-YCpBAC and 200 ng pTwist-OC43-N(CO) plasmids diluted in Opti-MEM I (Thermo Fisher, 31985070) and combined with 3.6 µL PEI MAX® (linear polyethylenimine (MW 40,000), 1 mg/mL in water (pH 7.4), 0.22 µm filtered; Polysciences, 24765) diluted in Opti-MEM I. Transfections were performed in serum-free DMEM for 4 h followed by a change of medium to 10% DMEM+Pen/Strep/Gln and the cells were left at 37°C. After 2 days, the transfected 293T cells were trypsinized and seeded into 10 cm dishes with BHK-21 cells and co-cultured for 4 days in 5% DMEM+Pen/Strep/Gln. Culture supernatants from these co-cultured cells were cleared by centrifugation (500 x *g* for 5 min) and used to infect monolayers of BHK-21 cells for 6 days (1^st^ passage). Culture supernatants from these 1^st^ passage infections were cleared by centrifugation and used to infect monolayers of BHK-21 cells for 2 days (2^nd^ passage). These 2^nd^ passage virus stocks were used to infect flasks of BHK-21 cells to generate virus stocks for subsequent experiments.

### Infections for time course experiments

293A cells were infected with OC43 (ATCC), OC43^YA^, or OC43-mClo^YA^ at an MOI of 0.1 in serum-free DMEM for 1 h at 37°C. The inocula were removed at 1 hour post-infection (hpi) and replaced with 2.5% DMEM+Pen/Strep/Gln prior to harvest at 1.5, 8, 16, 24, 32, or 48 hpi.

#### Titering

Viral supernatants from infected cells after the media change were collected and stored at −80°C prior to analysis by TCID50 assays as indicated above.

#### RNA samples for RT-qPCR

All medium was removed from the cells and the cells were lysed with 350 µL Buffer RLT Plus (QIAGEN). Cell lysates were stored at −80°C until processed for RNA isolations.

#### Cells lysates for western blotting

All medium was removed from the cells and the cells were gently washed with phosphate-buffered saline (PBS; Thermo Fisher, 10010023). After the PBS was removed, the cells were lysed with 2X Laemmli Buffer (12.5 mM Tris-HCl (pH 6.8), 4% SDS, 20% glycerol), and lysates were passed through QIAshredder columns (QIAGEN, 79654) for 2 min at 8000 x *g* to reduce viscosity. Samples were stored at −80°C until protein concentrations were measured and used for SDS-PAGE/western blotting.

#### Cells for flow cytometry

All medium was removed from the cells and the cells were gently washed with PBS prior to lifting the cells off the plates using 5 mM EDTA in PBS. The cells were pelleted at 500 x g for 3 min, after which the EDTA/PBS solution was removed, and the cells then fixed in 4% paraformaldehyde (PFA; Electron Microscopy Services, 15710) in PBS for 15 min at room temperature. The cells were pelleted at 2500 x *g* for 2 min, after which the PFA/PBS solution was removed, and the cells resuspended in PBS and stored at 4°C prior to staining for flow cytometry.

#### Cells for immunofluorescence

293A cells were seeded onto #1.5 round cover glass (VWR, MARI0117580) and infected as indicated above. All medium was removed from the cells and the cells were gently washed with PBS. The cells were then fixed in PFA in PBS for 15 min at room temperature. The fixation solution was removed, and the cells were stored in PBS at 4°C prior to staining for immunofluorescence.

### RT-qPCR

Total cell RNA was extracted from lysates prepared in Buffer RLT Plus using the RNeasy Plus Mini Kit (QIAGEN, 74136) according to the manufacturer’s instructions. Total RNA was converted to cDNA using the Maxima H Minus First Strand cDNA Synthesis Kit (Thermo Fisher, K1652) following the manufacturer’s “RT-qPCR-First Strand cDNA Synthesis” protocol using random hexamers and oligo (dT)_18_ primers with an extended 50°C synthesis step for 30 min. qPCRs were performed using Luna Universal qPCR Master Mix (NEB, M3003X) in 10 µL reactions with 200 nM primers (Table 5) and 1:200 diluted cDNA. One universal forward primer, 5’UTR (leader; Table 5), was used for all reactions. qPCRs were performed using CFX Connect Real-Time PCR Detection System (Bio-Rad) using the following two-step cycling conditions: 3 min at 95°C, 40 cycles of 10 s at 95°C and 30 s at 60°C followed by a 65-95°C melt curve. Analysis was performed using an efficiency-corrected 2^-ΔΔCq^ method (90).

**Table 5:**
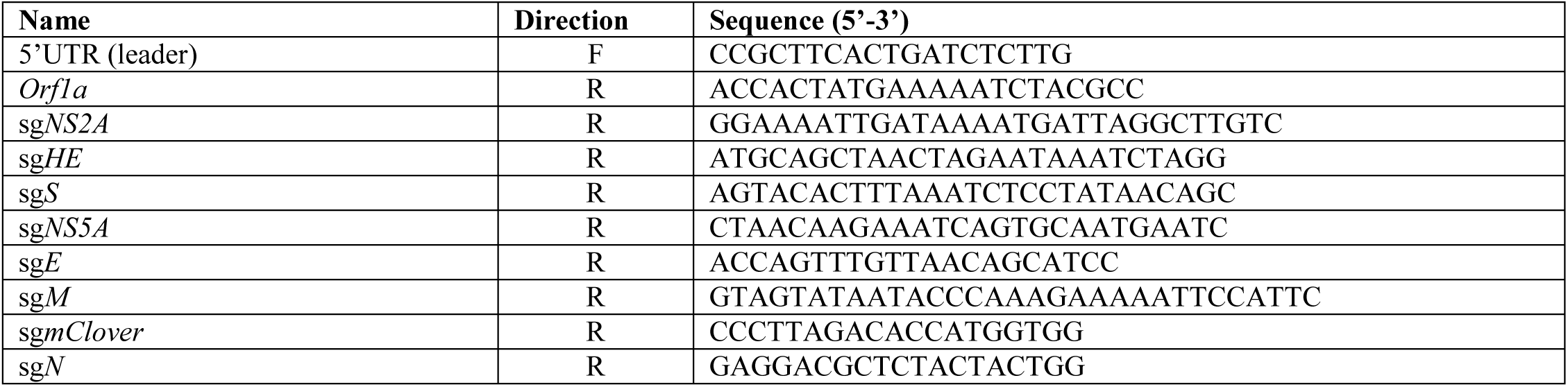
qPCR primers.

### SDS-PAGE and western blotting

Protein concentrations of lysates for SDS-PAGE were determined using the DC Protein Assay (Bio-Rad, 5000116) against a bovine serum albumin (BSA) standard curve and measured in a 96-well plate format at 750 nm using a Eon (BioTek) microplate spectrophotometer. Dithiothreitol (DTT; 108 mM final) and bromophenol blue (BPB; 0.001% final) were added to the protein lysates prior to loading. 5 µg of total protein was loaded into 6% or 12.5% acrylamide gels with Color Prestained Protein Standard, Broad Range (NEB, P7719S) and separated at 100 V. For samples used for PNGase F treatment, 4 µg of each lysate in 2X Laemmli buffer containing DTT/BPB was combined with GlycoBuffer 2 (1X final; NEB, P0704S) and NP-40 (1% final; NEB, P0704S) with or without 1 µL/reaction of PNGase F (NEB, P0704S) and incubated for 1 h at 37°C. prior to loading into 12.5% acrylamide gels. Proteins were transferred to polyvinylidene fluoride (PVDF) membranes using the Trans-Blot Turbo RTA Midi 0.2 µm PVDF Transfer Kit (Bio-Rad, 1704273) and a Trans-Blot Turbo Transfer System (Bio-Rad). Membranes were blocked (1 h, room temperature) and probed with primary (overnight, 4°C) and secondary antibodies (1 h, room temperature) in 4% BSA in TBST, except for blots for OC43-HE which were blocked in 4% skim milk in TBST. Primary antibodies used: Mouse anti-coronavirus, OC43 strain (OC43-N; Sigma-Aldrich, MAB9012, 1:2500); rabbit anti-GFP (mClover3-H2B; Cell Signaling, 2555, 1:1000); rabbit anti-OC43-S (CUSABIO, CSB-PA336163EA01HIY, 1:2000); rabbit anti-OC43-HE (M. Desforges, 1:500); and rabbit anti-β-Actin (HRP conjugate; Cell Signaling, 5125, 1:1000). Secondary antibodies used: Anti-rabbit IgG, HRP-linked (Cell Signaling, 7074, 1:5000) and anti-mouse IgG, HRP-linked (Cell Signaling, 7076, 1:5000). All blots were developed using Clarity ECL Western Blotting Substrate (Bio-Rad, 170-5061) and imaged using a ChemiDoc MP Imaging System (Bio-Rad).

### Flow cytometry

After fixation, the cells were pelleted at 2500 x *g* for 2 min and then permeabilized with 0.1% saponin (Sigma-Aldrich, 47036) in PBS (0.22 µm filtered) for 10 min at 4°C. The cells were stained in 0.1% saponin in PBS with mouse anti-OC43-N (MAB9012, 1:400) for 20 min at 4°C. The cells were washed twice with PBS with centrifugation at 2500 x *g* for 2 min. F(ab’)2 goat anti-mouse secondary antibody (Alexa Fluor 647; Thermo Fisher, A-21237, 1:500) was added in 0.1% saponin in PBS and incubated with the cells for 20 min at 4°C. After two washes with PBS (2500 x *g* for 2 min), the cells were resuspended in 1% BSA/1 mM EDTA in PBS (0.22 µm filtered) and stored at 4°C prior to analysis. Samples were analyzed using a CytoFLEX flow cytometer (Beckman Coulter) configured with 405 nm, 488 nm, and 638 nm lasers. Data analysis was performed using FCS Express v6.06.0040 (De Novo Software) to determine the percentage of mClover3-H2B and OC43-N (Alexa Fluor 647) positive single cells.

### Live cell fluorescence and immunofluorescence microscopy

For live cell imaging, the cell monolayers were visualized using an EVOS FL Cell Imaging System (Thermo Fisher) using a 20X objective. For immunofluorescence, cells grown and infected on cover glass were blocked and permeabilized in staining buffer (1% heat-inactivated human serum (Sigma-Aldrich, 4552), 0.1% Triton X-100 in PBS) for 1 h at room temperature, then stained with 1:500 dilution of mouse anti-dsRNA monoclonal antibody J2 (SCICONS, RNT-SCI-10010200) overnight at 4°C. The following day the cells were washed three times with PBS, then stained with chicken anti-mouse Alexa Fluor 647 (Thermo Fisher, A-21463) in staining buffer for 1 h at room temperature in the dark. The cells were washed three times with PBS prior to DNA staining with Hoescht 33342 (Thermo Fisher, 62249) for 5 min at room temperature. Following three final washes with PBS, the cover glasses were mounted on slides using ProLong Gold Antifade Mountant (Thermo Fisher, P36930). Z-stacks were imaged on a Zeiss LSM880 and processed into maximum intensity projections using Zen Black v1.0 (Zeiss).

### Reagents

All oligonucleotides >70 bp in length were PAGE-purified and purchased from Integrated DNA Technologies (IDT). Standard oligonucleotides (<40 bp) were purchased from IDT or Sigma-Aldrich. All restriction enzymes were purchased from NEB except for *Kfl*I (Thermo Fisher, FD2164). All plasmids were propagated in Stbl3 *E. coli* cells and purified using QIAGEN QIAprep Spin Miniprep (27104) or QIAfilter Plasmid Midi (12243) kits according to the manufacturer’s instructions.

### Data management and analysis

All data management was performed with Microsoft Excel for Microsoft 365. Nucleotide and protein sequences were analyzed, and plasmid maps generated using SnapGene v7.2.1 (Dotmatics). Graphing and statistical calculations were performed using GraphPad Prism for Windows v10.2.3. Figures were prepared using Affinity Designer v1.10.6.1665 (Serif).

## ACKNOWLEDGEMENTS

We thank the following colleagues for their assistance this work: Prashant J. Desai (Johns Hopkins School of Medicine) for providing *S. cerevisiae* strain VL6-48, pCC1BACYCp-URA, and technical support; Lauren M. Oldfield for technical support; Dr. David J. Kelvin (Dalhousie University) for providing HCoV-OC43 gRNA; Dr. Roy Duncan (Dalhousie University) for HCoV-OC43; Dr. Maya Shmulevitz (University of Alberta) for providing the T7-GFP plasmid; Dr. Jennifer A. Corcoran (University of Calgary) for providing the codon-optimized HCoV-OC43 N plasmid; Dr. M. Desforges (Institut National de la Recherche Scientifique) for the rabbit anti-HE anti-serum; Dr. Denys A. Khaperskyy (Dalhousie University) for the use of the EVOS Cell Imaging System; as well as the Cellular Molecular Digital Imaging and Flow Cytometry CORES facilities at Dalhousie University for their technical support. We also thank members of the Corcoran Lab, Dr. Che Colpitts (Queen’s University), and Dr. Guido van Marle (University of Calgary) for their thoughtful reviews of the manuscript. The following plasmids were gifts provided through Addgene: pKanCMV-mClover3-10aa-H2B (Michael Lin, http://n2t.net/addgene:74257, RRID:Addgene_74257); pKanCMV-mRuby3-10aa-H2B (Michael Lin, http://n2t.net/addgene:74258, RRID:Addgene_74258); mCardinal-C1 (Michael Davidson, http://n2t.net/addgene:54799, RRID:Addgene_54799); EBFP2-C1 (Michael Davidson, http://n2t.net/addgene:54665, RRID:Addgene_54665), and pGBW-m4134906 (pTwist-OC43-N(CO); Ginkgo Bioworks & Benjie Chen, http://n2t.net/addgene:151960, RRID:Addgene_151960). This work was supported by a grant awarded to C.M. from the Natural Sciences and Engineering Research Council (RGPIN/06800-2019). The funders had no role in study design, data collection and interpretation, or the decision to submit the work for publication.

## AUTHOR CONTRIBUTIONS

Conceptualization: B.A.D., E.S.P., and C.M.; Methodology: B.A.D., T.T., E.S.P., J.R.R., and C.M.; Formal Analysis: B.A.D.; Investigation: B.A.D., E.S.P., and T.T.; Resources: B.A.D. and T.T.; Writing – Original Draft: B.A.D. and C.M.; Writing –Review & Editing: B.A.D., T.T., E.S.P., J.R.R., and C.M.; Visualization: B.A.D.; Supervision: B.A.D. and C.M.; Project Administration: B.A.D. and C.M.; Funding Acquisition: C.M.

## REFERENCES

1. Saberi A, Gulyaeva AA, Brubacher JL, Newmark PA, Gorbalenya AE. 2018. A planarian nidovirus expands the limits of RNA genome size. PLOS Pathogens 14:e1007314.

2. Ferron F, Sama B, Decroly E, Canard B. 2021. The enzymes for genome size increase and maintenance of large (+)RNA viruses. Trends in Biochemical Sciences 46:866–877.

3. Malone B, Urakova N, Snijder EJ, Campbell EA. 2022. Structures and functions of coronavirus replication–transcription complexes and their relevance for SARS-CoV-2 drug design. Nat Rev Mol Cell Biol 23:21–39.

4. Yan S, Zhu Q, Hohl J, Dong A, Schlick T. 2023. Evolution of coronavirus frameshifting elements: Competing stem networks expalain conservation and variability. Proceedings of the National Academy of Sciences 120:e2221324120.

5. Jahirul Islam Md, Nawal Islam N, Siddik Alom Md, Kabir M, Halim MA. 2023. A review on structural, non-structural, and accesaory proteins of SARS-CoV-2: Highlighting drug target sites. Immunobiology 228:152302.

6. Sethna PB, Hung SL, Brian DA. 1989. Coronavirus subgenomic minus-strand RNAs and the potential for mRNA replicons. Proceedings of the National Academy of Sciences 86:5626– 5630.

7. Sawicki SG, Sawicki DL. 1990. Coronavirus transcription: subgenomic mouse hepatitis virus replicative intermediates function in RNA synthesis. Journal of Virology 64:1050– 1056.

8. Sethna PB, Hofmann MA, Brian DA. 1991. Minus-strand copies of replicating coronavirus mRNAs contain antileaders. Journal of Virology 65:320–325.

9. Koetzner CA, Parker MM, Ricard CS, Sturman LS, Masters PS. 1992. Repair and mutagenesis of the genome of a deletion mutant of the coronavirus mouse hepatitis virus by targeted RNA recombination. Journal of Virology 66:1841–1848.

10. de Haan Cornelis A. M., van Genne Linda, Stoop Jeroen N., Volders Haukeline, Rottier Peter J. M. 2003. Coronaviruses as Vectors: Position Dependence of Foreign Gene Expression. Journal of Virology 77:11312–11323.

11. van Beurden SJ, Berends AJ, Krämer-Kühl A, Spekreijse D, Chénard G, Philipp H-C, Mundt E, Rottier PJM, Verheije MH. 2017. A reverse genetics system for avian coronavirus infectious bronchitis virus based on targeted RNA recombination. Virology Journal 14:109.

12. Almazán F, González JM, Pénzes Z, Izeta A, Calvo E, Plana-Durán J, Enjuanes L. 2000. Engineering the largest RNA virus genome as an infectious bacterial artificial chromosome. Proceedings of the National Academy of Sciences 97:5516–5521.

13. Yount B, Curtis KM, Baric RS. 2000. Strategy for Systematic Assembly of Large RNA and DNA Genomes: Transmissible Gastroenteritis Virus Model. Journal of Virology 74:10600– 10611.

14. Thiel V, Herold J, Schelle B, Siddell SG. 2001. Infectious RNA transcribed in vitro from a cDNA copy of the human coronavirus genome cloned in vaccinia virus. Journal of General Virology 82:1273–1281.

15. Casais R, Thiel V, Siddell SG, Cavanagh D, Britton P. 2001. Reverse Genetics System for the Avian Coronavirus Infectious Bronchitis Virus. Journal of Virology 75:12359–12369.

16. Yount B, Denison MR, Weiss SR, Baric RS. 2002. Systematic Assembly of a Full-Length Infectious cDNA of Mouse Hepatitis Virus Strain A59. Journal of Virology 76:11065– 11078.

17. Coley SE, Lavi E, Sawicki SG, Fu L, Schelle B, Karl N, Siddell SG, Thiel V. 2005. Recombinant Mouse Hepatitis Virus Strain A59 from Cloned, Full-Length cDNA Replicates to High Titers In Vitro and Is Fully Pathogenic In Vivo. Journal of Virology 79:3097–3106.

18. St-Jean JR, Desforges M, Almazán F, Jacomy H, Enjuanes L, Talbot PJ. 2006. Recovery of a Neurovirulent Human Coronavirus OC43 from an Infectious cDNA Clone. Journal of Virology 80:3670–3674.

19. Almazán F, Dediego ML, Galán C, Escors D, Alvarez E, Ortego J, Sola I, Zuñiga S, Alonso S, Moreno JL, Nogales A, Capiscol C, Enjuanes L. 2006. Construction of a severe acute respiratory syndrome coronavirus infectious cDNA clone and a replicon to study coronavirus RNA synthesis. J Virol 80:10900–10906.

20. Tekes G, Hofmann-Lehmann R, Stallkamp I, Thiel V, Thiel H-J. 2008. Genome Organization and Reverse Genetic Analysis of a Type I Feline Coronavirus. Journal of Virology 82:1851–1859.

21. Donaldson EF, Yount B, Sims AC, Burkett S, Pickles RJ, Baric RS. 2008. Systematic Assembly of a Full-Length Infectious Clone of Human Coronavirus NL63. Journal of Virology 82:11948–11957.

22. Worm SHE van den, Eriksson KK, Zevenhoven JC, Weber F, Züst R, Kuri T, Dijkman R, Chang G, Siddell SG, Snijder EJ, Thiel V, Davidson AD. 2012. Reverse Genetics of SARS-Related Coronavirus Using Vaccinia Virus-Based Recombination. PLOS ONE 7:e32857.

23. Bálint Á, Farsang A, Zádori Z, Hornyák Á, Dencső L, Almazán F, Enjuanes L, Belák S. 2012. Molecular Characterization of Feline Infectious Peritonitis Virus Strain DF-2 and Studies of the Role of ORF3abc in Viral Cell Tropism. Journal of Virology 86:6258–6267.

24. Scobey T, Yount BL, Sims AC, Donaldson EF, Agnihothram SS, Menachery VD, Graham RL, Swanstrom J, Bove PF, Kim JD, Grego S, Randell SH, Baric RS. 2013. Reverse genetics with a full-length infectious cDNA of the Middle East respiratory syndrome coronavirus. Proceedings of the National Academy of Sciences 110:16157–16162.

25. Almazán F, DeDiego ML, Sola I, Zuñiga S, Nieto-Torres JL, Marquez-Jurado S, Andrés G, Enjuanes L. 2013. Engineering a Replication-Competent, Propagation-Defective Middle East Respiratory Syndrome Coronavirus as a Vaccine Candidate. mBio 4:10.1128/mbio.00650-13.

26. Jengarn J, Wongthida P, Wanasen N, Frantz PN, Wanitchang A, Jongkaewwattana A. 2015. Genetic manipulation of porcine epidemic diarrhoea virus recovered from a full-length infectious cDNA clone. Journal of General Virology 96:2206–2218.

27. Zhang M, Li W, Zhou P, Liu D, Luo R, Jongkaewwattana A, He Q. 2020. Genetic manipulation of porcine deltacoronavirus reveals insights into NS6 and NS7 functions: a novel strategy for vaccine design. Emerging Microbes & Infections 9:20–31.

28. Xie X, Muruato A, Lokugamage KG, Narayanan K, Zhang X, Zou J, Liu J, Schindewolf C, Bopp NE, Aguilar PV, Plante KS, Weaver SC, Makino S, LeDuc JW, Menachery VD, Shi P-Y. 2020. An Infectious cDNA Clone of SARS-CoV-2. Cell Host Microbe 27:841–848.e3.

29. Ye C, Chiem K, Park J-G, Oladunni F, Platt RN, Anderson T, Almazan F, de la Torre JC, Martinez-Sobrido L. 2020. Rescue of SARS-CoV-2 from a Single Bacterial Artificial Chromosome. mBio 11:e02168–20.

30. Rihn SJ, Merits A, Bakshi S, Turnbull ML, Wickenhagen A, Alexander AJT, Baillie C, Brennan B, Brown F, Brunker K, Bryden SR, Burness KA, Carmichael S, Cole SJ, Cowton VM, Davies P, Davis C, De Lorenzo G, Donald CL, Dorward M, Dunlop JI, Elliott M, Fares M, da Silva Filipe A, Freitas JR, Furnon W, Gestuveo RJ, Geyer A, Giesel D, Goldfarb DM, Goodman N, Gunson R, Hastie CJ, Herder V, Hughes J, Johnson C, Johnson N, Kohl A, Kerr K, Leech H, Lello LS, Li K, Lieber G, Liu X, Lingala R, Loney C, Mair D, McElwee MJ, McFarlane S, Nichols J, Nomikou K, Orr A, Orton RJ, Palmarini M, Parr YA, Pinto RM, Raggett S, Reid E, Robertson DL, Royle J, Cameron-Ruiz N, Shepherd JG, Smollett K, Stewart DG, Stewart M, Sugrue E, Szemiel AM, Taggart A, Thomson EC, Tong L, Torrie LS, Toth R, Varjak M, Wang S, Wilkinson SG, Wyatt PG, Zusinaite E, Alessi DR, Patel AH, Zaid A, Wilson SJ, Mahalingam S. 2021. A plasmid DNA-launched SARS-CoV-2 reverse genetics system and coronavirus toolkit for COVID-19 research. PLoS Biol 19:e3001091.

31. Chiem K, Morales Vasquez D, Park J-G, Platt RN, Anderson T, Walter MR, Kobie JJ, Ye C, Martinez-Sobrido L. 2021. Generation and Characterization of Recombinant SARS-CoV-2 Expressing Reporter Genes. Journal of Virology 95:10.1128/jvi.02209-20.

32. Diefenbacher MV, Baric TJ, Martinez DR, Baric RS, Catanzaro NJ, Sheahan TP. 2024. A nano-luciferase expressing human coronavirus OC43 for countermeasure development. Virus Research 339:199286.

33. Terada Y, Kuroda Y, Morikawa S, Matsuura Y, Maeda K, Kamitani W. 2019. Establishment of a Virulent Full-Length cDNA Clone for Type I Feline Coronavirus Strain C3663. J Virol 93:e01208–19.

34. Herrmann A, Jungnickl D, Cordsmeier A, Peter AS, Überla K, Ensser A. 2021. Cloning of a Passage-Free SARS-CoV-2 Genome and Mutagenesis Using Red Recombination. Int J Mol Sci 22:10188.

35. Lee JY, Bae S, Myoung J. 2019. Generation of Full-Length Infectious cDNA Clones of Middle East Respiratory Syndrome Coronavirus 29:999–1007.

36. Torii S, Ono C, Suzuki R, Morioka Y, Anzai I, Fauzyah Y, Maeda Y, Kamitani W, Fukuhara T, Matsuura Y. 2021. Establishment of a reverse genetics system for SARS-CoV-2 using circular polymerase extension reaction. Cell Reports 35:109014.

37. Amarilla AA, Sng JDJ, Parry R, Deerain JM, Potter JR, Setoh YX, Rawle DJ, Le TT, Modhiran N, Wang X, Peng NYG, Torres FJ, Pyke A, Harrison JJ, Freney ME, Liang B, McMillan CLD, Cheung STM, Guevara DJDC, Hardy JM, Bettington M, Muller DA, Coulibaly F, Moore F, Hall RA, Young PR, Mackenzie JM, Hobson-Peters J, Suhrbier A, Watterson D, Khromykh AA. 2021. A versatile reverse genetics platform for SARS-CoV-2 and other positive-strand RNA viruses. Nat Commun 12:3431.

38. Kim BK, Choi W-S, Jeong JH, Oh S, Park J-H, Yun YS, Min SC, Kang DH, Kim E-G, Ryu H, Kim HK, Baek YH, Choi YK, Song M-S. 2023. A Rapid Method for Generating Infectious SARS-CoV-2 and Variants Using Mutagenesis and Circular Polymerase Extension Cloning. Microbiology Spectrum 11:e03385–22.

39. Liu G, Gack MU. 2023. An optimized circular polymerase extension reaction-based method for functional analysis of SARS-CoV-2. Virol J 20:63.

40. Hu Z, López-Muñoz AD, Kosik I, Li T, Callahan V, Brooks K, Yee DS, Holly J, Santos JJS, Castro Brant A, Johnson RF, Takeda K, Zheng Z-M, Brenchley JM, Yewdell JW, Fox JM. 2024. Recombinant OC43 SARS-CoV-2 spike replacement virus: An improved BSL-2 proxy virus for SARS-CoV-2 neutralization assays. Proceedings of the National Academy of Sciences 121:e2310421121.

41. Taha TY, Chen IP, Hayashi JM, Tabata T, Walcott K, Kimmerly GR, Syed AM, Ciling A, Suryawanshi RK, Martin HS, Bach BH, Tsou C-L, Montano M, Khalid MM, Sreekumar BK, Renuka Kumar G, Wyman S, Doudna JA, Ott M. 2023. Rapid assembly of SARS-CoV-2 genomes reveals attenuation of the Omicron BA.1 variant through NSP6. Nat Commun 14:2308.

42. Mélade J, Piorkowski G, Touret F, Fourié T, Driouich J, Cochin M, Bouzidi HS, Coutard B, Nougairède A, de Lamballerie X. 2022. A simple reverse genetics method to generate recombinant coronaviruses. EMBO reports 23:e53820.

43. Kipfer ET, Hauser D, Lett MJ, Otte F, Urda L, Zhang Y, Lang CM, Chami M, Mittelholzer C, Klimkait T. Rapid cloning-free mutagenesis of new SARS-CoV-2 variants using a novel reverse genetics platform. eLife 12:RP89035.

44. Yang Y-L, Liang Q-Z, Xu S-Y, Mazing E, Xu G-H, Peng L, Qin P, Wang B, Huang Y-W. 2019. Characterization of a novel bat-HKU2-like swine enteric alphacoronavirus (SeACoV) infection in cultured cells and development of a SeACoV infectious clone. Virology 536:110–118.

45. Thi Nhu Thao T, Labroussaa F, Ebert N, V’kovski P, Stalder H, Portmann J, Kelly J, Steiner S, Holwerda M, Kratzel A, Gultom M, Schmied K, Laloli L, Hüsser L, Wider M, Pfaender S, Hirt D, Cippà V, Crespo-Pomar S, Schröder S, Muth D, Niemeyer D, Corman VM, Müller MA, Drosten C, Dijkman R, Jores J, Thiel V. 2020. Rapid reconstruction of SARS-CoV-2 using a synthetic genomics platform. Nature 582:561–565.

46. Cao H, Gu H, Kang H, Jia H. 2023. Development of a rapid reverse genetics system for feline coronavirus based on TAR cloning in yeast. Front Microbiol 14.

47. Kouprina N, Larionov V. 2016. Transformation-associated recombination (TAR) cloning for genomics studies and synthetic biology. Chromosoma 125:621–632.

48. Vashee S, Stockwell TB, Alperovich N, Denisova EA, Gibson DG, Cady KC, Miller K, Kannan K, Malouli D, Crawford LB, Voorhies AA, Bruening E, Caposio P, Früh K. 2017. Cloning, Assembly, and Modification of the Primary Human Cytomegalovirus Isolate Toledo by Yeast-Based Transformation-Associated Recombination. mSphere 2.

49. Oldfield LM, Grzesik P, Voorhies AA, Alperovich N, MacMath D, Najera CD, Chandra DS, Prasad S, Noskov VN, Montague MG, Friedman RM, Desai PJ, Vashee S. 2017. Genome-wide engineering of an infectious clone of herpes simplex virus type 1 using synthetic genomics assembly methods. Proc Natl Acad Sci USA 114:E8885–E8894.

50. Knickmann J, Staliunaite L, Puhach O, Ostermann E, Günther T, Nichols J, Jarvis MA, Voigt S, Grundhoff A, Davison AJ, Brune W. 2024. A simple method for rapid cloning of complete herpesvirus genomes. Cell Rep Methods 4:100696.

51. Zhou Y, Li C, Ren C, Hu J, Song C, Wang X, Li Y. 2022. One-Step Assembly of a Porcine Epidemic Diarrhea Virus Infectious cDNA Clone by Homologous Recombination in Yeast: Rapid Manipulation of Viral Genome With CRISPR/Cas9 Gene-Editing Technology. Front Microbiol 13.

52. Li C, Chen X, Zhou Y, Hu J, Wang X, Li Y. 2022. Novel reverse genetics of genotype I and III Japanese encephalitis viruses assembled through transformation associated recombination in yeast: The reporter viruses expressing a green fluorescent protein for the antiviral screening assay. Antiviral Research 197:105233.

53. Bajar BT, Wang ES, Lam AJ, Kim BB, Jacobs CL, Howe ES, Davidson MW, Lin MZ, Chu J. 2016. Improving brightness and photostability of green and red fluorescent proteins for live cell imaging and FRET reporting. Sci Rep 6:20889.

54. Shen L, Yang Y, Ye F, Liu G, Desforges M, Talbot PJ, Tan W. 2016. Safe and Sensitive Antiviral Screening Platform Based on Recombinant Human Coronavirus OC43 Expressing the Luciferase Reporter Gene. Antimicrob Agents Chemother 60:5492–5503.

55. Larionov V, Kouprina N, Solomon G, Barrett JC, Resnick MA. 1997. Direct isolation of human BRCA2 gene by transformation-associated recombination in yeast. Proceedings of the National Academy of Sciences 94:7384–7387.

56. Gibson DG, Benders GA, Andrews-Pfannkoch C, Denisova EA, Baden-Tillson H, Zaveri J, Stockwell TB, Brownley A, Thomas DW, Algire MA, Merryman C, Young L, Noskov VN, Glass JI, Venter JC, Hutchison CA 3rd, Smith HO. 2008. Complete chemical synthesis, assembly, and cloning of a Mycoplasma genitalium genome. Science 319:1215–1220.

57. Gibson DG, Glass JI, Lartigue C, Noskov VN, Chuang R-Y, Algire MA, Benders GA, Montague MG, Ma L, Moodie MM, Merryman C, Vashee S, Krishnakumar R, Assad-Garcia N, Andrews-Pfannkoch C, Denisova EA, Young L, Qi Z-Q, Segall-Shapiro TH, Calvey CH, Parmar PP, Hutchison CA, Smith HO, Venter JC. 2010. Creation of a Bacterial Cell Controlled by a Chemically Synthesized Genome. Science 329:52–56.

58. Zhao Y, Coelho C, Hughes AL, Lazar-Stefanita L, Yang S, Brooks AN, Walker RSK, Zhang W, Lauer S, Hernandez C, Cai J, Mitchell LA, Agmon N, Shen Y, Sall J, Fanfani V, Jalan A, Rivera J, Liang F-X, Bader JS, Stracquadanio G, Steinmetz LM, Cai Y, Boeke JD. 2023. Debugging and consolidating multiple synthetic chromosomes reveals combinatorial genetic interactions. Cell 186:5220–5236.e16.

59. Sharmeen L, Kuo MY, Dinter-Gottlieb G, Taylor J. 1988. Antigenomic RNA of human hepatitis delta virus can undergo self-cleavage. Journal of Virology 62:2674–2679.

60. Kuo MY, Sharmeen L, Dinter-Gottlieb G, Taylor J. 1988. Characterization of self-cleaving RNA sequences on the genome and antigenome of human hepatitis delta virus. Journal of Virology 62:4439–4444.

61. Yount B, Curtis KM, Fritz EA, Hensley LE, Jahrling PB, Prentice E, Denison MR, Geisbert TW, Baric RS. 2003. Reverse genetics with a full-length infectious cDNA of severe acute respiratory syndrome coronavirus. Proceedings of the National Academy of Sciences 100:12995–13000.

62. St-Jean JR, Desforges M, Talbot PJ. 2006. Genetic Evolution of Human Coronavirus OC43 in Neural Cell Culture, p. 499–502. In Perlman, S, Holmes, KV (eds.), The Nidoviruses. Springer US, Boston, MA.

63. Siragam V, Maltseva M, Castonguay N, Galipeau Y, Srinivasan MM, Soto JH, Dankar S, Langlois M-A. 2024. Seasonal human coronaviruses OC43, 229E, and NL63 induce cell surface modulation of entry receptors and display host cell-specific viral replication kinetics. Microbiology Spectrum 12:e04220–23.

64. Fung TS, Liu DX. 2018. Post-Translational Modifications of Coronavirus Proteins: Roles and Function. Future Virol 13:405–430.

65. Lutomski CA, El-Baba TJ, Bolla JR, Robinson CV. 2021. Multiple Roles of SARS-CoV-2 N Protein Facilitated by Proteoform-Specific Interactions with RNA, Host Proteins, and Convalescent Antibodies. JACS Au 1:1147–1157.

66. Meyer B, Chiaravalli J, Gellenoncourt S, Brownridge P, Bryne DP, Daly LA, Grauslys A, Walter M, Agou F, Chakrabarti LA, Craik CS, Eyers CE, Eyers PA, Gambin Y, Jones AR, Sierecki E, Verdin E, Vignuzzi M, Emmott E. 2021. Characterising proteolysis during SARS-CoV-2 infection identifies viral cleavage sites and cellular targets with therapeutic potential. Nat Commun 12:5553.

67. Chu H, Hou Y, Yang D, Wen L, Shuai H, Yoon C, Shi J, Chai Y, Yuen TT-T, Hu B, Li C, Zhao X, Wang Y, Huang X, Lee KS, Luo C, Cai J-P, Poon VK-M, Chan CC-S, Zhang AJ, Yuan S, Sit K-Y, Foo DC-C, Au W-K, Wong KK-Y, Zhou J, Kok K-H, Jin D-Y, Chan JF- W, Yuen K-Y. 2022. Coronaviruses exploit a host cysteine-aspartic protease for replication. Nature 609:785–792.

68. Millet JK, Whittaker GR. 2015. Host cell proteases: Critical determinants of coronavirus tropism and pathogenesis. Virus Research 202:120–134.

69. Desforges M, Desjardins J, Zhang C, Talbot PJ. 2013. The Acetyl-Esterase Activity of the Hemagglutinin-Esterase Protein of Human Coronavirus OC43 Strongly Enhances the Production of Infectious Virus. J Virol 87:3097–3107.

70. Hurdiss DL, Drulyte I, Lang Y, Shamorkina TM, Pronker MF, van Kuppeveld FJM, Snijder J, de Groot RJ. 2020. Cryo-EM structure of coronavirus-HKU1 haemagglutinin esterase reveals architectural changes arising from prolonged circulation in humans. Nat Commun 11:4646.

71. Zhang R, Wang K, Ping X, Yu W, Qian Z, Xiong S, Sun B. 2015. The ns12.9 Accessory Protein of Human Coronavirus OC43 Is a Viroporin Involved in Virion Morphogenesis and Pathogenesis. J Virol 89:11383–11395.

72. The transcriptional and translational landscape of HCoV-OC43 infection | bioRxiv. https://www.biorxiv.org/content/10.1101/2024.01.20.576440v1. Retrieved 14 August 2024.

73. Konings DA, Bredenbeek PJ, Noten JF, Hogeweg P, Spaan WJ. 1988. Differential premature termination of transcription as a proposed mechanism for the regulation of coronavirus gene expression. Nucleic Acids Res 16:10849–10860.

74. van Marle G, Luytjes W, van der Most RG, van der Straaten T, Spaan WJM. 1995. Regulation of coronavirus mRNA transcription. Journal of Virology 69:7851–7856.

75. Owen DR, Allerton CMN, Anderson AS, Aschenbrenner L, Avery M, Berritt S, Boras B, Cardin RD, Carlo A, Coffman KJ, Dantonio A, Di L, Eng H, Ferre R, Gajiwala KS, Gibson SA, Greasley SE, Hurst BL, Kadar EP, Kalgutkar AS, Lee JC, Lee J, Liu W, Mason SW, Noell S, Novak JJ, Obach RS, Ogilvie K, Patel NC, Pettersson M, Rai DK, Reese MR, Sammons MF, Sathish JG, Singh RSP, Steppan CM, Stewart AE, Tuttle JB, Updyke L, Verhoest PR, Wei L, Yang Q, Zhu Y. 2021. An oral SARS-CoV-2 Mpro inhibitor clinical candidate for the treatment of COVID-19. Science 374:1586–1593.

76. Weil T, Lawrenz J, Seidel A, Münch J, Müller JA. 2022. Immunodetection assays for the quantification of seasonal common cold coronaviruses OC43, NL63, or 229E infection confirm nirmatrelvir as broad coronavirus inhibitor. Antiviral Research 203:105343.

77. Li J, Wang Y, Solanki K, Atre R, Lavrijsen M, Pan Q, Baig MS, Li P. 2023. Nirmatrelvir exerts distinct antiviral potency against different human coronaviruses. Antiviral Research 211:105555.

78. Makino S, Joo M. 1993. Effect of intergenic consensus sequence flanking sequences on coronavirus transcription. Journal of Virology 67:3304–3311.

79. Edridge AWD, Kaczorowska J, Hoste ACR, Bakker M, Klein M, Loens K, Jebbink MF, Matser A, Kinsella CM, Rueda P, Ieven M, Goossens H, Prins M, Sastre P, Deijs M, van der Hoek L. 2020. Seasonal coronavirus protective immunity is short-lasting. Nature Medicine 26:1691–1693.

80. Shah MM, Winn A, Dahl RM, Kniss KL, Silk BJ, Killerby ME. 2022. Seasonality of Common Human Coronaviruses, United States, 2014-2021(1). Emerg Infect Dis 28:1970–1976.

81. Kistler KE, Bedford T. 2021. Evidence for adaptive evolution in the receptor-binding domain of seasonal coronaviruses OC43 and 229e. eLife 10:e64509.

82. Dugas M, Grote-Westrick T, Merle U, Fontenay M, Kremer AE, Hanses F, Vollenberg R, Lorentzen E, Tiwari-Heckler S, Duchemin J, Ellouze S, Vetter M, Fürst J, Schuster P, Brix T, Denkinger CM, Müller-Tidow C, Schmidt H, Tepasse P-R, Kühn J. 2021. Lack of antibodies against seasonal coronavirus OC43 nucleocapsid protein identifies patients at risk of critical COVID-19. Journal of Clinical Virology 139:104847.

83. Abela Irene A., Schwarzmüller Magdalena, Ulyte Agne, Radtke Thomas, Haile Sarah R., Ammann Priska, Raineri Alessia, Rueegg Sonja, Epp Selina, Berger Christoph, Böni Jürg, Manrique Amapola, Audigé Annette, Huber Michael, Schreiber Peter W., Scheier Thomas, Fehr Jan, Weber Jacqueline, Rusert Peter, Günthard Huldrych F., Kouyos Roger D., Puhan Milo A., Kriemler Susi, Trkola Alexandra, Pasin Chloé. 2024. Cross-protective HCoV immunity reduces symptom development during SARS-CoV-2 infection. mBio 15:e02722–23.

84. Meslin FX, Kaplan MM, Koprowski H, World Health Organization. 1996. Laboratory techniques in rabies4th ed. Monograph 23:445–459.

85. Chu J, Haynes RD, Corbel SY, Li P, González-González E, Burg JS, Ataie NJ, Lam AJ, Cranfill PJ, Baird MA, Davidson MW, Ng H-L, Garcia KC, Contag CH, Shen K, Blau HM, Lin MZ. 2014. Non-invasive intravital imaging of cellular differentiation with a bright red-excitable fluorescent protein. Nat Methods 11:572–578.

86. Lartigue C, Vashee S, Algire MA, Chuang R-Y, Benders GA, Ma L, Noskov VN, Denisova EA, Gibson DG, Assad-Garcia N, Alperovich N, Thomas DW, Merryman C, Hutchison CA, Smith HO, Venter JC, Glass JI. 2009. Creating Bacterial Strains from Genomes That Have Been Cloned and Engineered in Yeast. Science 325:1693–1696.

87. Kouprina N, Larionov V. 2008. Selective isolation of genomic loci from complex genomes by transformation-associated recombination cloning in the yeast Saccharomyces cerevisiae. Nat Protoc 3:371–377.

88. Vashee S, Stockwell TB, Alperovich N, Denisova EA, Gibson DG, Cady KC, Miller K, Kannan K, Malouli D, Crawford LB, Voorhies AA, Bruening E, Caposio P, Früh K. 2017. Cloning, Assembly, and Modification of the Primary Human Cytomegalovirus Isolate Toledo by Yeast-Based Transformation-Associated Recombination. mSphere 2:e00331–17.

89. Nadai Y, Eyzaguirre LM, Constantine NT, Sill AM, Cleghorn F, Blattner WA, Carr JK. 2008. Protocol for Nearly Full-Length Sequencing of HIV-1 RNA from Plasma. PLoS One 3:e1420.

90. Damgaard MV, Treebak JT. 2022. Protocol for qPCR analysis that corrects for cDNA amplification efficiency. STAR Protocols 3:101515.

